# Molecular dating of the emergence of anaerobic rumen fungi and the impact of laterally acquired genes

**DOI:** 10.1101/401869

**Authors:** Yan Wang, Noha Youssef, M.B. Couger, Radwa Hanafy, Mostafa Elshahed, Jason E. Stajich

## Abstract

The anaerobic gut fungi (AGF) or Neocallimastigomycota inhabit the rumen and alimentary tract of herbivorous mammals, where they play an important role in the degradation of plant fiber. Comparative genomic and phylogenomic analysis of the AGF has long been hampered by their fastidious growth pattern as well as their large and AT-biased genomes. We sequenced 21 AGF transcriptomes and combined them with 5 available genome sequences of AGF taxa to explore their evolutionary relationships, time their divergence, and characterize patterns of gene gain/loss associated with their evolution. We estimate that the most recent common ancestor of the AGF diverged 66 (±10) million years ago, a timeframe that coincides with the evolution of grasses (Poaceae), as well as the mammalian transition from insectivory to herbivory. The concordance of these independently estimated ages of AGF evolution, grasses evolution, and mammalian transition to herbivory suggest that AGF have been important in shaping the success of mammalian herbivory transition by improving the efficiency of energy acquisition from recalcitrant plant materials. Comparative genomics identified multiple lineage-specific genes and protein domains in the AGF, two of which were acquired from an animal host (galectin) and rumen gut bacteria (carbohydrate-binding domain) via horizontal gene transfer (HGT). Four of the bacterial derived “Cthe_2159” genes in AGF genomes also encode eukaryotic Pfam domains (“Atrophin-1”, “eIF-3_zeta”, “Nop14”, and “TPH”) indicating possible gene fusion events after the acquisition of “Cthe_2159” domain. A third AGF domain, plant-like polysaccharide lyase N-terminal domain (“Rhamnogal_lyase”), represents the first report from fungi that potentially aids AGF to degrade pectin. Analysis of genomic and transcriptomic sequences confirmed the presence and expression of these lineage-specific genes in nearly all AGF clades supporting the hypothesis that these laterally acquired and novel genes in fungi are likely functional. These genetic elements may contribute to the exceptional abilities of AGF to degrade plant biomass and enable metabolism of the rumen microbes and animal hosts.

## Introduction

Diverse microbes inhabit the digestive tract of ruminant mammals and contribute to degradation of ingested plant fibers, a process that liberates nutrients for their hosts. Large scale genomic and metagenomic sequencing of rumen microbes have produced hundreds of novel bacterial genomes enabling discovery of plant biomass degrading enzymes and patterns of genomic evolution (Seshadri et al., 2018; Stewart et al., 2018). However, eukaryotic members of the rumen microbial community have been less intensely studied (Haitjema et al., 2017; Youssef et al., 2013). Members of the phylum Neocallimastigomycota (anaerobic gut fungi or AGF) are important members of the rumen and hindgut of a wide range of herbivorous mammals and reptiles (Gruninger et al., 2014). To survive in this anoxic and prokaryotes-dominated environment, extant AGF members have undergone multiple structures and metabolic adaptations, including the loss of the mitochondria, a gain of a hydrogenosome, the loss of respiratory capacities, and a substitution of ergosterol with tetrahymanol in the cell membrane (Yarlett et al., 1986). Importantly, all known AGF taxa have a remarkably efficient plant biomass degradation machinery, which may be critical for competing with other microbes for resources and establishing growth in the herbivorous gut. Such capacity is reflected in the possession of an impressive arsenal of plant biomass degradation enzymes and the production of the cellulosomes—extracellular structures that harbor multiple enzymes bound to scaffoldins (Haitjema et al., 2017). These metabolic and structural adaptations improve survivability, fitness, and competitiveness of the AGF in the herbivorous gut, but the genetic and evolutionary origins of these changes remain largely undescribed (Solomon et al., 2016; Youssef et al., 2013). Previous genomic investigations of the AGF have identified a massive number of carbohydrate active enzymes coded by genes with foreign origins presumably from multiple lineages of bacteria through Horizontal Gene Transfer (HGT) events independently (Haitjema et al., 2017; Solomon et al., 2016; Youssef et al., 2013). In fact, HGT examples from bacteria to fungi have been documented extensively (Chaib De Mares et al., 2014; Dhillon et al., 2015; Gardiner et al., 2012; Pombert et al., 2012). However, HGT elements in fungi that have been transferred from other eukaryotes are still rare with only a few described cases from animals (Wang et al., 2016), oomycetes (Sun et al., 2011), or plants (Richards et al., 2009). The rumen is an intriguing context to explore patterns of HGT, where degradative enzymes break down sorts of cells liberating DNA and RNA molecules. Competing organisms can find advantage by acquiring foreign genes that operate efficiently in an anaerobic environment to obtain nutrients from recalcitrant plant fibers or to recognize other microbes.

The Neocallimastigomycota are classified within the Chytridiomycota (chytrid) fungi, which share the trait of a flagellated zoospore stage (James et al., 2006a, 2006b; Spatafora et al., 2017; Stajich, 2017). Efforts to resolve the phylogenetic relationship of AGF and their sister lineages using ribosomal markers have yielded conflicting topologies (Liggenstoffer et al., 2010; Wang et al., 2017). Multilocus phylogeny or phylogenomics has not yet been applied to evaluate their evolutionary relationships and to estimate the divergence time of the AGF. Using genomes and transcriptomes from 26 different AGF taxa (Table 1) covering seven out of the ten recognized genera, we reconstructed a robust phylogenomic tree of the AGF and estimated their divergence time. We compared the genomes or transcriptomes of AGF and their non-rumen associated relatives in Chytridiomycota to identify unique and shared genome contents. This study examined the relatively recent divergence of the AGF clade and revealed a concordance of the divergence time of the Neocallimastigomycota fungi with both the mammalian hosts transition to herbivory and the diversification events of the forage grasses. As the AGF are well known for their exceptional efficiency at plant biomass degradation, we also explored the diverse genetic components of these fungi. We discovered two potential HGT elements that were found unique to the AGF, which are predicted to have originated from animals or bacteria. Examination of the family of bacterial transferred genes revealed multiple intron insertion events that occurred after the HGT acquisition process, which are present in all five AGF genomes. Comparative analyses of these genes suggest the intron insert events were related to intragenic duplication of coding sequences. In addition, a novel plant polysaccharide lyase was revealed from both AGF genomes and transcriptomes that has never been reported from any known fungal genomes or genetic studies. The evolutionary genomic investigation of these rumen inhabiting fungi provides perspective on the concordant timing of their divergence with the ecological niche they inhabit and the potential role of HGT in accumulation of lineage-specific processes that may contribute to their unique biology.

**Table 1.**
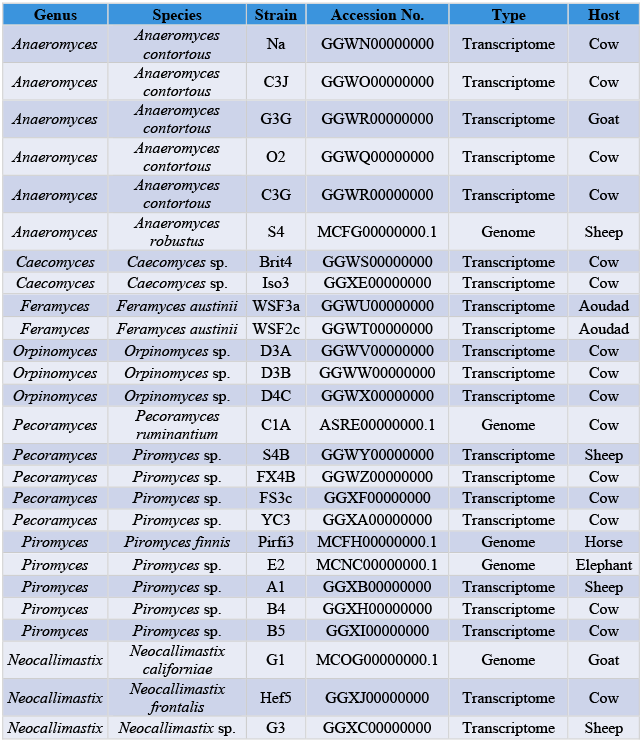
Information for the AGF strains included in this study.

## Results

### Divergence time estimation and phylogenomic relationship of Neocallimastigomycota

Phylogenomic analysis placed the 26 AGF taxa into a single monophyletic clade with strong support of Bayesian posterior probability (1.0/1.0) and maximum likelihood bootstrap value (100%) (Figure 1 and Figure S1). All AGF genera *(Anaeromyces, Caecomyces, Feramyces, Neocallimastix*, *Orpinomyces*, *Pecoramyces*, and *Piromyces*) included in this study formed individual monophyletic clades that were also supported by both Bayesian (Figure 1) and maximum likelihood analyses (Figure S1). A conflict in the tree topology between the two phylogenetic reconstructions is the placement of the *Caecomyces* clade. This lineage is sister to the rest of the Neocallimastigomycota in the maximum likelihood tree (Figure S1), while the *Caecomyces* position is swapped with *Piromyces* in the Bayesian phylogeny (Figure 1). This is likely due to short internode distances, which suggest a rapid radiation of the ancestors of the two genera. The relative short bar of the highest-probability density (HPD) on the node of the AGF clade (Figure 1) suggests the integrative natural history of this group of fungi and the outperforming resolving power of the genome-wide data in the molecular dating analyses.

**Figure 1.**
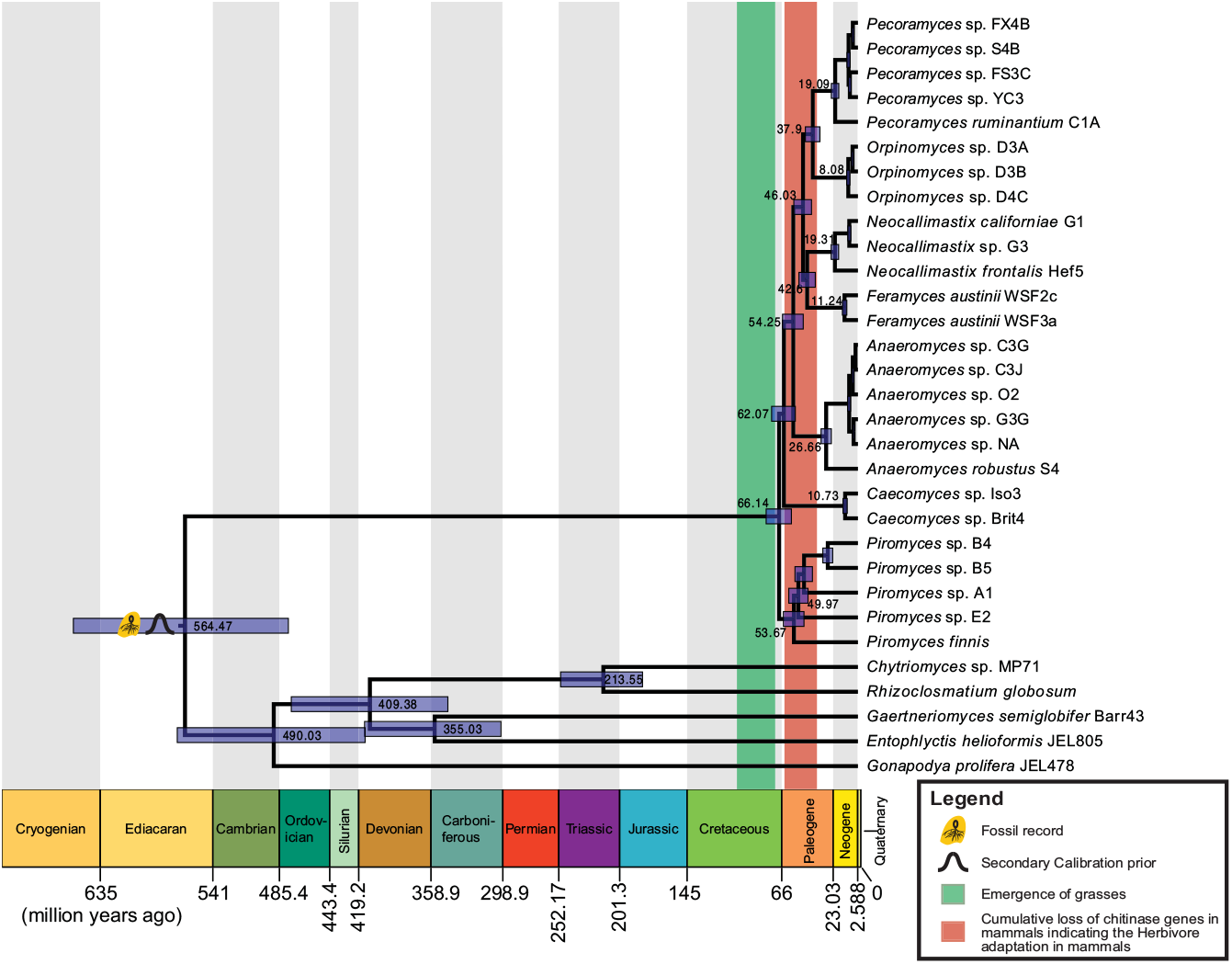
Bayesian phylogenomic Maximum Clade Credibility tree of Neocallimastigomycota with divergence time estimation. All clades are fully supported by Bayesian posterior probabilities (BPP). For clarity, mean ages and 95% highest-probability density ranges (blue bars) are denoted on the nodes above the rank of genus.

The divergence time of the Neocallimastigomycota clade is estimated at the Cretaceous/Paleogene (K/Pg) boundary 66 (±10) Mya (Figure 1). The chronogram (Figure 1) displays a long branch leading to the emergence of the AGF clade, which extends from the end of Ediacaran (~564 Mya) to the K/Pg boundary (~66 Mya). This suggests that the extant members of AGF did not emerge until recently and then rapidly radiated into separate clades in the Paleogene. The estimated time frame for AGF divergence broadly coincides with the age of the grasses (70-95 Mya), previously estimated using molecular (nuclear and chloroplast) markers, and calibrated using fossils from pollen and dinosaur coprolite as well as the breakup time of the Gondwana (Bremer, 2002; Christin et al., 2014; Gaut, 2002; Prasad et al., 2005; Vicentini et al., 2008). In addition, this inferred AGF divergence time also coincides with a major diet change of placental mammals: the transition from a primarily insectivorous to a herbivorous and omnivorous lifestyles. The loss of chitinase genes diversity, estimated to occurred from the Cretaceous/Paleogene (K/Pg) boundary (66 Mya) to the mid of Paleogene (34 Mya) (Figure 1), is widely seen as a consequence of such transition (Emerling et al., 2018). Collectively, these overlapping estimates suggest that the evolution of the symbiotic association between herbivorous mammals and rumen fungi is tightly linked with the evolution of forage grasses and mammalian dietary transitions within a 66-95 Mya timeframe. The exact chronology of these three divergence or transition events cannot be accurately determined partially due to the intervals of the estimates (Figure 1). However, the dates inferred from phylogenetic analyses are consistent with the hypothesis that rumen fungi have played important roles in the diet transition of some mammals to acquire nutrition from forage grasses.

### Genome-wide comparison of protein domains and homologous genes

Comparative genomic analysis between AGF and their non-rumen associated chytrid relatives (Figure 2) identified 40 Pfam domains that are unique to the AGF, representing 0.67% of the total number of Pfams (5,980) in the AGF pan-genome-transcriptome (Table S1 and Figure 2b). The predicted functions of these domains include anaerobic ribonucleotide reductase (“NRDD”), metal transport and binding (“FeoA”, “FeoB_C”), carbohydrate binding (e.g., “CBM_10”, “CBM-like”, “Cthe_2159”), atypical protein kinase (“CotH”), and glycoside hydrolase (e.g., “Glyco_hydro_6”, “Glyco_hydro_11”) (Table S1 and Figure 2b). In addition to these 40 unique AGF domains, many additional Pfams were also enriched in the AGF. Such domains mediate polysaccharide degradation and monosaccharide fermentations (Figure 2c), including “Chitin_binding_1”, “CBM_1”, “Cellulase”, “Glyco_hydro_10”, “Gly_radical”, “RicinB_lectin_2”, “Esterase”, and “Polysacc_deac_1” domains. Further, our analysis also identified 106 Pfam domains that are not present in AGF genomes and transcriptomes but found in sister Chytridiomycota. Most of these missing domains are related to oxidation reactions on cytochromes and mitochondria, instead, they possess specialized organelle called hydrogenosome conducting metabolism in the anaerobic condition (Yarlett et al., 1986) (Table S1 and Figure 2d). In addition, domains involved in the biosynthesis of nicotinic acid, uric acid, purine catabolism, photolyase, and pathways of ureidoglycolate and kynurenine are also found to be absent in AGF species. Similar patterns were also identified in the comparison of homologous genes (Figure S2).

**Figure 2.**
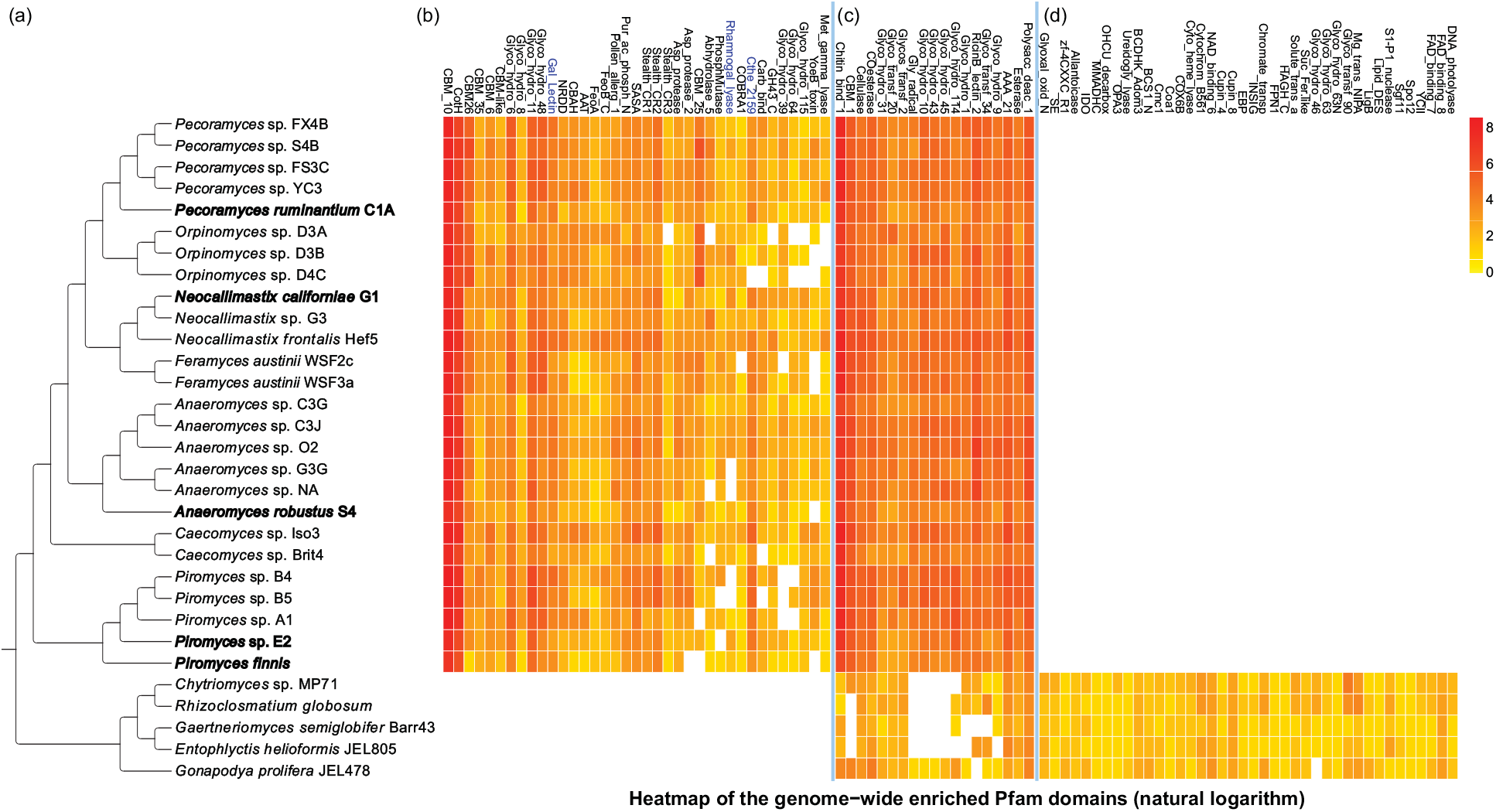
Cladogram and heatmap enrichment of the Pfam domains between Neocallimastigomycota and Chytridiomycota. (a) Cladogram showing the phylogenetic relationship of the compared taxa (Neocallimastigomycota genomes are in bold, transcriptomes in standard type); (b) Heatmap plot of natural logarithm of the domain copy numbers showing the ones uniquely gained in Neocallimastigomycota (suggested HGT elements are in blue); (c) Pfam domains highly enriched in Neocallimastigomycota; and (d) Pfam domains absent in Neocallimastigomycota (presented domains are partial; see Table S1 for the full list).

A permissive criterion, allowing some missing copies, found a total of 2,728 gene families shared between AGF and chytrids. We discovered that 1,709 additional gene families are shared among AGF genomes (each gene presents in at least 21 out of the total 26 taxa) but absent in other chytrids, while another 367 families are missing in AGF members but present in the other chytrid lineages.

### Genomic interactions within the rumen of mammalian herbivores

We focused on three Pfam domains (“Cthe_2159”, “Gal_Lectin”, and “Rhamnogal_lyase”) that are unique to the Neocallimastigomycota and previously not observed in fungal genomes. Phylogenetic analyses support a horizontal transfer of “Cthe_2159” from rumen bacteria into AGF followed by potential gene fusion to deliver eukaryotic specific functions. Similarly, analysis of “Gal_Lectin” domain copies in AGF suggests they were acquired from animal donor lineages. Similarity search of AGF “Rhamnogal_lyase” domain finds most similar copies in plant genomes and phylogenetic analysis indicates the AGF polysaccharide lyase domain is distinct and not orthologous to related enzymes in other fungi.

#### A bacteria-like biomass-binding and putatively polysaccharide lyase domain (“Cthe_2159”)

The “Cthe_2159” domain was originally characterized as a polysaccharide lyase-like protein in the thermophilic and biomass-degrading bacterium *Clostridium thermocellum* (Close et al., 2014). “Cthe_2159” are beta-helix proteins with the ability to bind celluloses and acid-sugars (polygalacturonic acid, a major component of the pectin) and homologs are primarily found in archaeal and bacterial genomes. Notably, a total 583 copies of the “Cthe_2159” domain were identified in 5 genomes and 21 transcriptomes of AGF taxa, but reduced to a set of 126 clusters based on overall protein similarity (>90%) due to redundancy in transcriptome assemblies. This domain is absent in all other eukaryotic genomes examined in this study (Figure 3 and Table 2). A phylogenetic tree of “Cthe_2159” homologs identified from Archaea, Bacteria, and AGF suggest that the AGF “Cthe_2159” domains were acquired from bacteria through HGT (Figure 3). The likely donor was a gram-positive Firmicute *(Clostridiales)* (Maximum Likelihood bootstrap value 98%) and the closest protein copies of “Cthe_2159” domains are encoded in the *Oribacterium sinus, Oribacterium* sp., and *Hungatella hathewayi* genomes (Figure 3). Members of the order Clostridiales are integral members of the rumen microbiome. Four of these AGF “Cthe_2159” domain containing genes also encode eukaryotic Pfam protein domains (“Atrophin-1”, “eIF-3_zeta”, “Nop14”, and “TPH”) at the 3’ position of the “Cthe_2159” domain. We hypothesize these domains are the result of fusion after the acquisition of “Cthe_2159” domain. The functions of these additional domains include initiation of the eukaryotic translation, maturation of 18S rRNA, production of 40S ribosome, and meiosis-specific activities (Figure 4a). Approximately 30% of these AGF “Cthe_2159” gene models possess between 1 and 2 introns but there is limited spliced transcript evidence to provide confidence in the gene structures, so the apparent intron gains could be artifacts of genome assembly or annotation (Supplementary results) (Haitjema et al., 2017; Youssef et al., 2013).

**Figure 3.**
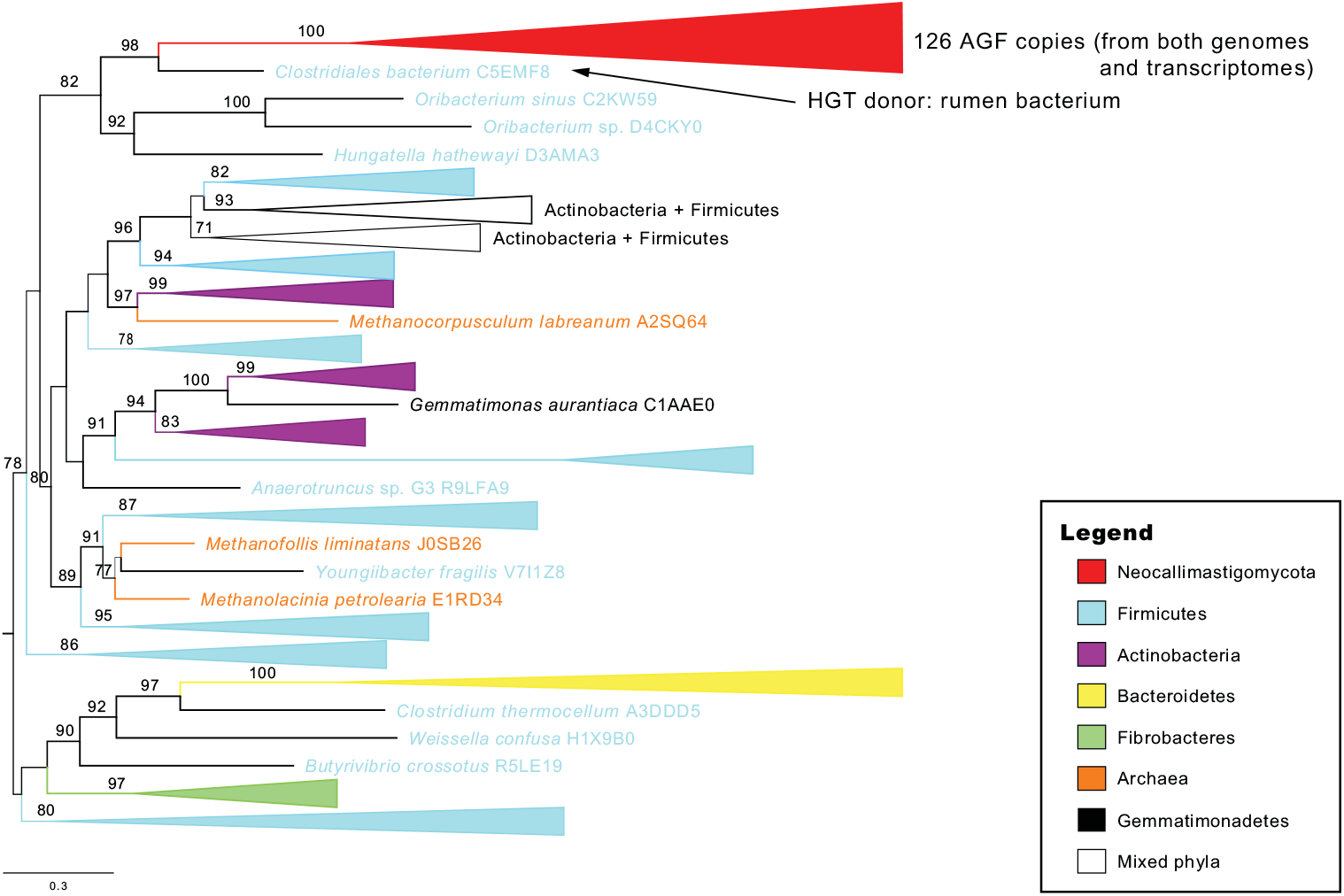
Mid-point rooted phylogenetic tree of the “Cthe_2159” domain. All 126 Neocallimastigomycota (AGF) copies (copies that >90% identities have been removed) form a single clade (red) indicating the HGT donor, *Clostridiales bacterium* C5EMF8 (an obligate rumen bacterium), with strong support of maximum likelihood bootstrap (98/100). Included bacterial lineages were assigned different colors according to their phylogenetic classification (see legend for detailed information; the complete tree with all tip information is in the Figure S3).

**Figure 4.**
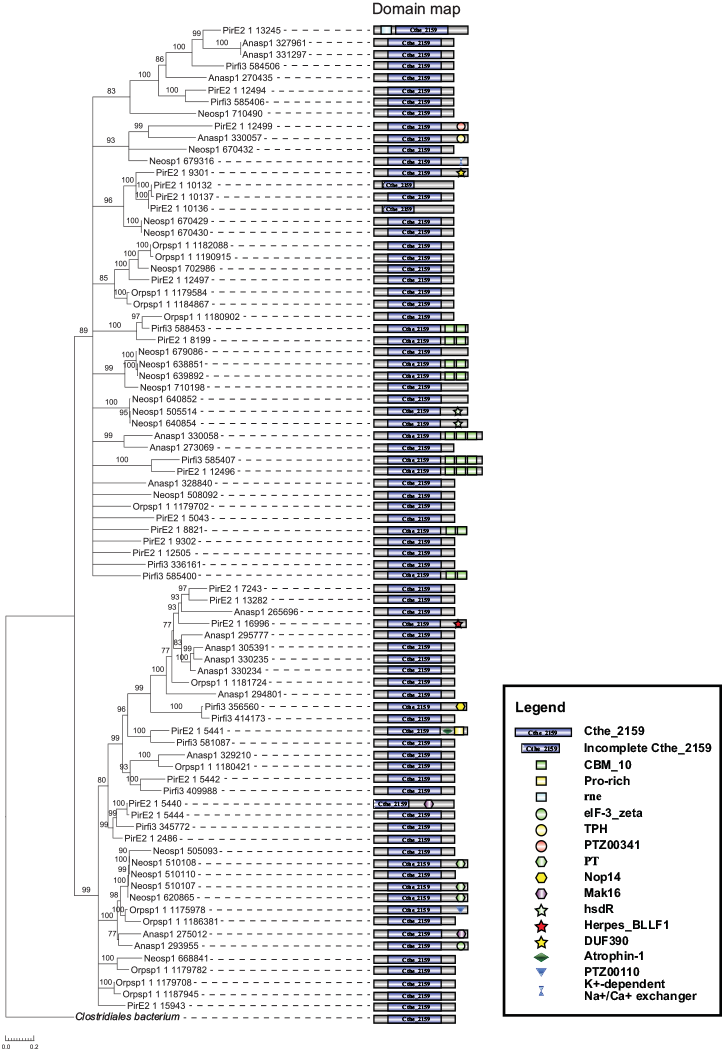
Phylogenetic tree of the 83 “Cthe_2159” domains identified in five AGF genomes based on protein sequences (rooted with the closest related bacterial homolog). Domain maps on the right shows the conserved domains produced by the “Cthe_2159” containing genes.

**Table 2.**
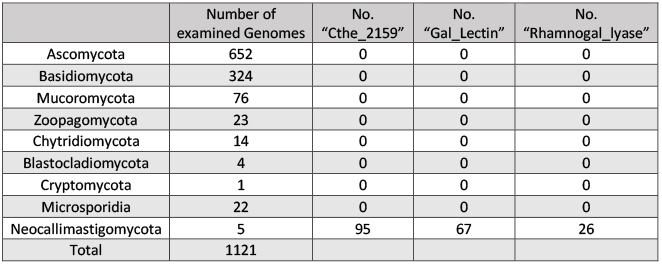
Distribution of the three studied domains in the fungal kingdom.

|  | Number of<br>examined Genomes | No.<br>“Cthe_2159” | No.<br>“Gal_Lectin” | No.<br>“Rhamnogal_lyase” |
| --- | --- | --- | --- | --- |
| Ascomycota | 652 | 0 | 0 | 0 |
| Basidiomycota | 324 | 0 | 0 | 0 |
| Mucoromycota | 76 | 0 | 0 | 0 |
| Zoopagomycota | 23 | 0 | 0 | 0 |
| Chytridiomycota | 14 | 0 | 0 | 0 |
| Blastocladiomycota | 4 | 0 | 0 | 0 |
| Cryptomycota | 1 | 0 | 0 | 0 |
| Microsporidia | 22 | 0 | 0 | 0 |
| Neocallimastigomycota | 5 | 95 | 67 | 26 |
| Total | 1121 |  |  |  |

#### An animal-like galactose binding lectin domain (“Gal_Lectin”)

“Gal_Lectin” domains were found in AGF genomes universally and absent in all other examined chytrid and fungal genomes (Table 2). Phylogenetic analysis recovered a monophyletic AGF “Gal_Lectin” clade which was not placed as a sister clade to the animals as expected for a fungal gene. Instead, it was embedded within the animal homologs in the tree and allies with one subgroup, polycystin-1 (PC-1 protein) (Figure 5a). The three separate animal subclades contain protein members that harbor the “Gal_Lectin” domain but with dissimilar sequence information and annotated functions (Figure 5). The genomes of ruminant hosts (e.g., horse, sheep) of the AGF fungi also encode three gene families containing the “Gal_Lectin” domain, which can be observed in each of the animal subclades (Figure 5). The proteins in the animal subclade 1 were annotated as the PC-1, which are homologues to the human “polycystic kidney disease” *(PKD1)* genes. The members of the animal subclade 2 were searched by BLAST for the the “adhesion G-protein coupled receptor L1/3” (ADGRL1). The animal subclade 3 contains homologs of the “EVA-1 protein”, most of which contain two adjacent copies of the “Gal_Lectin” domain. The three subgroups of animal “Gal_Lectin” domains are also flanked by disparate Pfam domains (Figure 5b). The gene phylogeny suggests an animal PC-1 protein as the likely donor lineage for the AGF “Gal_Lectin” gene (Figure 5a), based on its closest sister relationship. In addition, the AGF proteins also contain a Pfam “Glyco_transf_34” domain (Figure 5b) which is absent in all animal homologs of the “Gal_Lectin” containing genes suggesting its involvement in fungus specific activities in the rumen.

**Figure 5.**
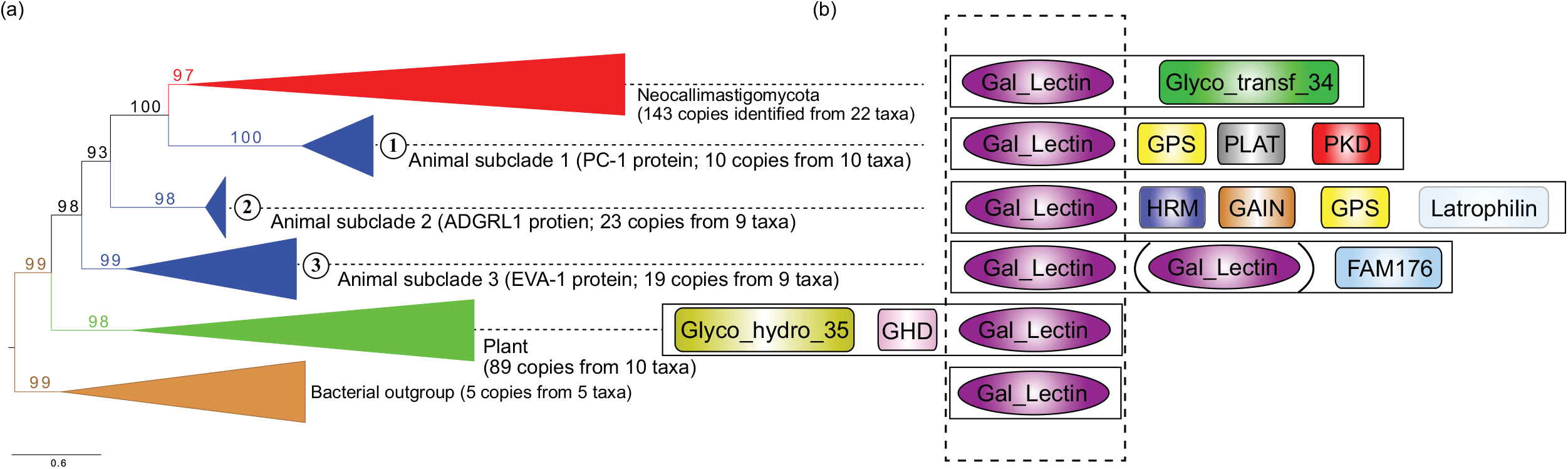
Phylogenetic tree of the animal-like “Gal-Lectin” domain identified in Neocallimastigomycota. (a) Collapsed phylogenetic tree based on protein sequences (rooted with the bacterial outgroup), including clades of Neocallimastigomycota (red), Animals (blue; three clades are labeled as 1-3), plants (green), and bacteria (brown) (A complete tree with all tip information is in the Figure S4); (b) Schematic diagrams showing the “Gal_Lectin” and other conserved domains on the same protein in each clade individually (dotted box highlights the aligned region used to produce the phylogenetic tree).

#### A novel fungal rhamnogalacturonate lyase (“Rhamnogal_lyase”) in AGF

In plants, the rhamnogalacturonate lyases are involved in the fruit ripening-related process, cell-wall modification, and lateral-root and root-hair formation (Molina-Hidalgo et al., 2013; Ponniah et al., 2017). The Pfam database classifies two types of domains for rhamnogalactoside degrading activity: “Rhamnogal_lyase” and “RhgB_N”. They are both N-terminal catalytic domains associated with the rhamnogalacturonan lyase protein (polysaccharide lyase family 4, PL4) and flanked persistently by the group of “fn3_3” and “CBM-like” domains, with the particular function to degrade the rhamnogalacturonan I (RG-I) backbone of pectin. The “Rhamnogal_lyase” domain exists in members of plants and plant-pathogenic bacteria (e.g. *Erwinia chrysanthemi),* whereas the “RhgB_N” domain has a wider distribution and can be found in bacteria, fungi, and oomycetes (Finn et al., 2016). Sequence similarity searches using the “Rhamnogal_lyase” domain against various protein sequence databases (e.g., EnsEMBL, Mycocosm, Pfam) returned no homolog in any other fungi (except the AGF members), which indicates that this domain is unique to AGF, plants, and bacteria. On the other hand, the “RhgB_N” domain is widely shared by Dikarya fungi, Oomycetes, and bacteria. Although “RhgB_N” and “Rhamnogal_lyase” domains are distantly related according to the sequence similarity (24% between the copies of the *Aspergillus nidulans* and *An. robustus),* they presumably share a common origin due to that they both physically located on the N-terminal region of the PL4 proteins and they have resembling functions to degrade the pectin RG-I region. The phylogenetic tree shows that although AGF “Rhamnogal_lyase” domains are more closely related to the plant homologs than to the clades of fungi and oomycetes, these AGF rhamnogalacturonate lyases likely have evolved a specific function in fungi (Figure 3). The presence of the “Rhamnogal_lyase” domain in the rumen-associated fungi suggests that the AGF may support an ability to soften, modify, and degrade the plant pectin within the anaerobic rumen in a related but different way from plants.

## Discussion

Microbial diversity of ruminants is a research hotspot for development of bioenergy tools (Bryant, 1959; Marvin-Sikkema et al., 1994; Seshadri et al., 2018). The AGF fungi are an important but understudied component of the ruminant microbiome and their obligate anaerobic and relatively A-T rich genomes have limited the initial genomic resources for the group. In this study, we produced the most phylogenetically broad transcriptome sampling of the Neocallimastigomycota fungi to date to support phylogenomic and comparative analyses. Our results contribute new insights into the natural history and dynamic evolution of these cryptic ruminant gut fungi. The reconstructed phylogenomic species tree resolved previously unanswered questions about the evolutionary relationships of the members of the AGF. In addition, we provide the first estimation of the divergence time of AGF taxa, 66 (±10) Mya (Figure 1), which is in remarkable concordance with the divergence of the forage Poaceae grasses (70-95 Mya) and dietary shifts in mammalian lineages (34-66 Mya) from insectivore to herbivore and omnivore. Grass evolution enabled the herbivory transition, and this diet adaptation drove an increase in developmental and morphological complexity of the digestive tract, compartmentalization, and the development of dedicated anaerobic fermentation chambers (e.g., rumen and caecum) in the herbivorous alimentary tract to improve biomass degradation efficiency (Hackmann and Spain, 2010). This transition to plant-based (or plant-exclusive) diets required additional partnership with microbes since mammals lack cellulolytic and hemi-cellulolytic enzymes necessary to liberate sugars for absorption (Gruninger et al., 2014). In addition, the genome content comparisons help illustrate and predict new biological roles AGF play in the mammalian herbivore guts. The long branch that leads to the emergence of the Neocallimastigomycota clade indicates the distinctiveness of the extant group of obligate symbiotic fungi in the mammalian herbivores and implies the existence of undiscovered although possibly extinct relatives of the Neocallimastigomycota and Chytridiomycota (Figure 1). Future environmental and metagenome sequence exploration of anaerobic environments testing for presence of these types of fungi may provide new observations that support their existence.

Our analyses identified multiple instances of Pfam domain gains (n=40) and losses (n=106) within the Neocallimastigomycota clade (Figure 2 and Table S1). We identified three AGF lineage specific protein domains which are absent from all other examined fungal genomes (Table 2). Phylogenetic analyses support the hypothesis that they were acquired via HGT or other noncanonical events. Phylogenetic analyses of “Cthe_2159” and “Gal_Lectin” indicate the domains were separately transferred from the rumen bacteria and animal hosts horizontally (Figures 3 and 5). The gains of these domains highlight how HGT has contributed to broaden the lignocellulolytic capacities (through “Cthe_2159”) of the AGF and potentially increase their abilities for cell recognition (through “Gal_Lectin”) within the rumen. The presence of four eukaryotic Pfam domains fused with these bacteria-originated “Cthe_2159” genes in AGF suggests they are truly eukaryotic and encoded in the fungal genomes (Figure 4) and not a contamination artifact. Studies of intron gains and losses in fungal lineages have suggested the ancestor was intron rich, an observation that is supported by intron rich chytrid genomes (Csuros et al., 2011; Nielsen et al., 2004; Stajich et al., 2007). Although introns are present in gene models of several “Cthe_2159” copies found in all available AGF genomes, and many are flanked with duplicated coding sequences (in *Piromyces* sp. E2), we are not able to confidently conclude these models experienced recent intron-insertion events as there is little support of spliced mRNA transcripts originating from these loci (Supplementary results).

The “Cthe_2159” is a newly described protein family that bind cellulosic and pectic substrates in the anaerobic and thermophilic bacterium *Clostridium thermocellum* (Close et al., 2014). The crystal structure of the “Cthe_2159” suggests that it is a polysaccharide lyase family with similarity with pectate lyases in the PL9 family. Similarly, “Rhamnogal_lyase” domains are primarily function in the facilitation of cell wall modification in plants (Molina-Hidalgo et al., 2013). Phytopathogenic bacteria can utilize their prokaryotic versions of the domain to disorganize plant tissues to support the invasion (Laatu and Condemine, 2003). Although we cannot locate the original donor lineages of the AGF “Rhamnogal_lyase” domains (Figure 6), their gain is a key synapomorphy of the extant AGF taxa and may contribute to the ability of these fungi to access polysaccharides in plant cell walls. Both “Cthe_2159” and “Rhamnogal_lyase” (PL4 family) domains function in pectin binding or degradation activities, and the possession both suggests that AGF may have evolved abilities to deconstruct pectin with an exceptional efficiency that distinguish themselves from other fungi (Table 2 and Figure 2). Pectin is abundant in primary cell walls and the middle lamella in both dicotyledonous plants (making up 20-35% dry weight) and grasses (2-10%) serving as a protection of plant cells from degrading enzymes produced by animals (Salem et al., 2017; Vogel, 2008; Voragen et al., 2009; Xiao and Anderson, 2013). Removal of pectin can effectively increase the surface of exposed plant cell wall, and thus improve the accessibility of other polysaccharides (cellulose and hemicellulose) masked by pectin (Pakarinen et al., 2012). Both of the “Cthe_2159” and “Rhamnogal_lyase” proteins may have contributed to the high efficiency of the AGF biomass degradability by uncoupling the pectin that glues cells together, increasing the exposed surface areas, and thus allowing diverse polysaccharide enzymes to work on plant cells simultaneously in the rumen. These protein domains could account for the superior performance of AGF to weaken forage fibers and release polysaccharides (Borneman et al., 1989; Nagpal et al., 2009). The AGF may benefit or depend on these acquired domains in their capacity as primary degraders of ingested forage (Haitjema et al., 2014).

**Figure 6.**
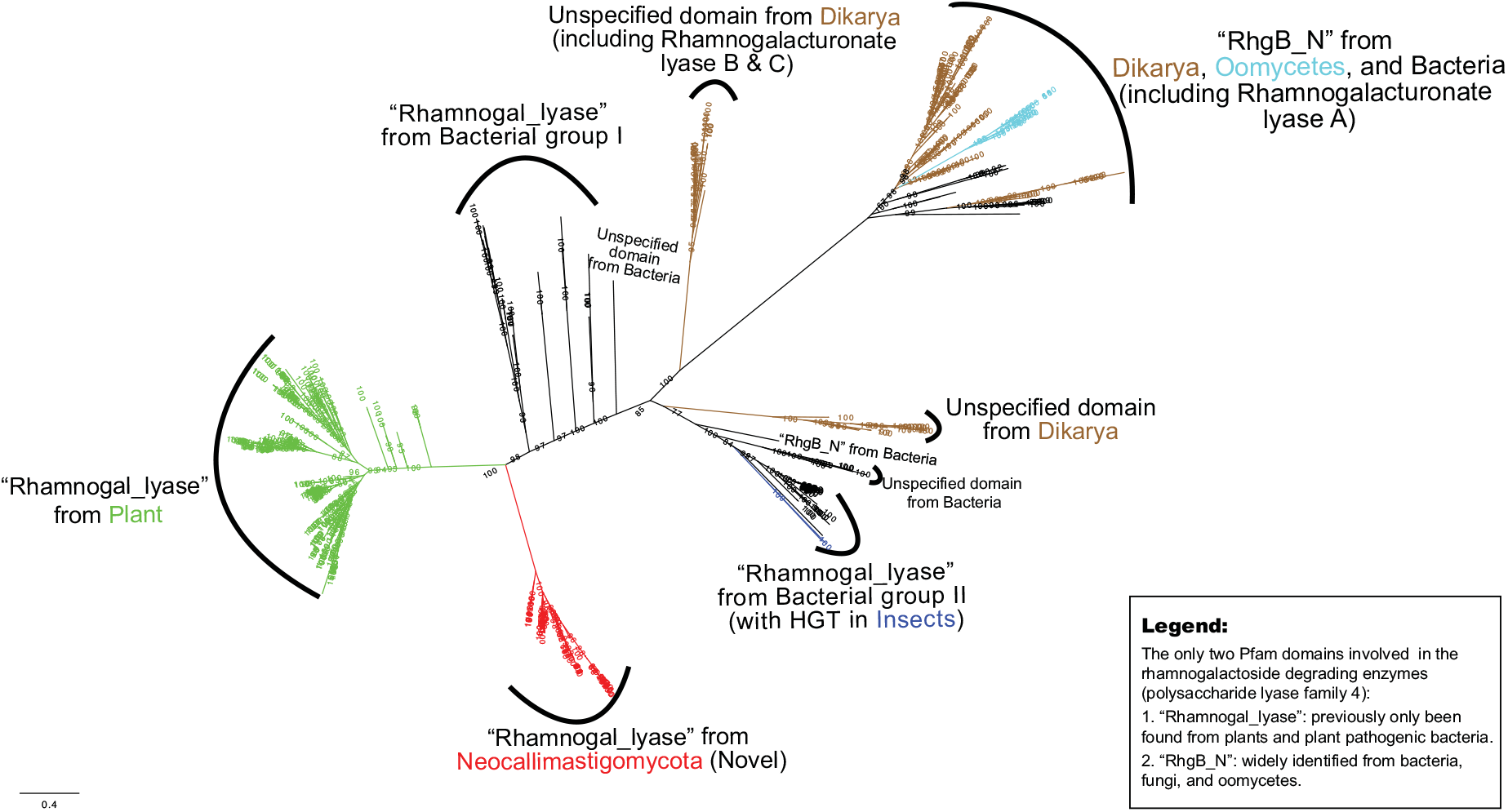
Radial phylogenetic tree of the “Rhamnogal_lyase” domain encoded by the Neocallimastigomycota (red). Plant copies are colored in green and other homologous fungal genes are colored in brown. Oomycetes are in cyan and animal copies only known in the mountain pine beetles *Dendroctonus ponderosae* are in blue. Bacterial branches remain in black. The tree also included homologs of “RhgB_N” and “Rhamnogalacturonan lyase A, B, and C”. Domain names are suggested using NCBI’s conserved domain search tool (cutoff 1E^-5^) with unaligned FASTA sequences (refer to the Figure S6 for a tree with detailed information).

The “Gal_Lectin” domain bears the phylogenetic hallmark of being acquired from an animal donor. Animals use galactose-binding lectins to recognize foreign entities (García-Maldonado et al., 2017) and participate in anti-microbial defenses (Low et al., 2010; Uhlenbruck and Steinhausen, 1977). Our results suggest that the “Gal_Lectin” domains in AGF are homologous and closely related to animal PC-1 proteins (Figure 5a), which are transmembrane proteins functioning in cell recognition (Hughes et al., 1995; Weston et al., 2003). *In vitro,* PC-1 shows binding ability to carbohydrate matrices and collagens type I, II, and IV (Weston et al., 2001). As such, we postulate that the acquisition of the animal-like “Gal_Lectin” domain contributes to the AGF abilities of cell-cell recognition and interaction with other microbes in the rumen. Syntenic relationship of the coding genes shows that the AGF “Gal_Lectin” domains are flanked by the “Glyco_transf_34” domain, which lacks of homologs in any other animals (Figure 5b and Figure S5). The AGF-equipped “Glyco_transf_34” belongs to the galactosyl transferase GMA12/MNN10 family and may help catalyze the transfer of the sugar moieties in cooperating with the adjacent “Gal_Lectin” domain. Our investigation found that HGT has contributed to the AGF genome evolution with donors from both prokaryotes and eukaryotes. HGT may have helped these fungi to acquire new functions and to thrive in the anaerobic gut as a key member of the microbial community degrading plant materials in animal hosts.

Other than the arsenal of diverse enzyme profiles, the AGF have also been known to use rhizoids and holdfasts to physically aid the fungal body to penetrate into the plant material deeply, which is superior than other rumen microorganisms in terms of efficiency (Berlemont, 2017; Gruninger et al., 2014). Our study provides evidence that the rumen fungi are able to and have actively acquire functional domains from the animal hosts and co-existing anaerobic bacteria in the rumen. These exotic genetic elements encoded in Neocallimastigomycota genomes may contribute to their distinctive function comprised of unique genomic assets comparing to their free-living relatives. The long branch leading to the recent radiation of Neocallimastigomycota (Figure 1) also suggests distinct evolutionary trajectory from the sister Chytridiomycota lineages. Living as gut-dwellers in the strict anaerobic gut environment for over 66 million years, AGF have undergone reductive evolution on the mitochondria and eventually transformed it to a new organelle—hydrosome (Marvin-Sikkema et al., 1994; Youssef et al., 2013). Their ecological roles of AGF in such an extreme environment also endow their exceptional ability for plant degradation. The AGF use both physical (deconstruction of lignocelluloses) and biological (depolymerization) mechanisms before the fermentation of plant polysaccharides. These steps require diverse enzymes capable of breaking chemical bonds in carbohydrates including cellulases, hemicellulases, ligninases, and pectinases (Brandt et al., 2013). These all drive the synapomorphic and autapomorphic characters discovered in AGF. Currently few close relatives have been found and none cultured which subtend the long branch. Environmental DNA investigations of extreme environment that may be a suitable niche of those Neocallimastigomycota-like microbes may reveal potential relatives (James et al., 2000). For example, a recent metagenomic survey from costal marine sediments suggests that some operational taxonomic units (OTUs) could be assigned to Neocallimastigomycota using 28S rRNA marker (Picard, 2017). Sampling of deep sea habitats and marine mammalian herbivores could provide future discoveries of biodiversity and evolutionary importance for understanding the evolutionary trajectory of the Neocallimastigomycota.

## Materials and Methods

### Transcriptome and genome datasets

We generated the transcriptomes of 21 strains of Neocallimastigomycota fungi from cow, sheep, horse, and goat feces, and rumen fluid of fistulated cows in the Stillwater, OK area (Murphy et al., n.d.) (Table 1). These strains were maintained under anaerobic conditions using the modified Hungate method as described previously (Balch and Wolfe, 1976; Bryant, 1972; Hanafy et al., 2017; Hungate and Macy, 1973). Total volume of RNA was harvested from the growing fungal strains and processed for transcriptomics sequencing, which was performed using an external commercial service provided by Novogene (Beijing, China). The RNAseq data were assembled into *de novo* transcript assemblies using Trinity (v2.6.6), followed by TransDecoder (v5.0.2) to predict ORFs (Haas et al., 2013). The generated proteomes and corresponding coding sequences were used as input to phylogenomic and comparative genomic analyses. The five published Neocallimastigomycota genome sequences were obtained from JGI MycoCosm database (Grigoriev et al., 2014; Spatafora, 2011). These are *Anaeromyces robustus* S4, *Neocallimastix californiae* G1, *Pecoramyces ruminantium* C1A (synonym *Orpinomyces* sp.), *Piromyces finnis (v3.0),* and *Piromyces* sp. E2 (Haitjema et al., 2017; Youssef et al., 2013). Five outgroup Chytridiomycota taxa with sequenced genomes were chosen. These are *Chytriomyces* sp. MP 71, *Entophlyctis helioformis* JEL805, *Gaertneriomyces semiglobifer* Barr 43, *Gonapodya prolifera* JEL478, and *Rhizoclosmatium globosum* JEL800 (Chang et al., 2015; Mondo et al., 2017). Assembled transcriptomes, raw Illumina read sequences, and isolates metadata are deposited in the GenBank with the BioProject ID PRJNA489922. All accession numbers are listed in the Table 1.

### Phylogenomics and divergence time estimation

A set of 434 highly conserved and generally single-copy protein coding genes in fungi and animal and plant outgroups (DOI: 10.5281/zenodo.1413687) were used for phylogenomic analyses in the PHYling pipeline (DOI: 10.5281/zenodo.1257002). Profile-Hidden-Markov-Models of these markers were searched in the chytrid predicted protein sequences using HMMER3 (v3.1b2). A total of 426 (out of 434) conserved orthologous markers were identified with hmmsearch (cutoff=1E^-10^) in the 26 Neocallimastigomycota and 5 Chytridiomycota. The identified protein sequence homologs in each species, for each phylogenetic marker, were aligned with hmmalign to the marker profile-HMM. The protein alignments were also back translation into codon alignments guided by the protein alignment using the tool bp_mrtrans.pl (Stajich et al., 2002). The protein and coding sequences of the markers were concatenated into a super-alignment with 426 partitions defined by each gene marker. The 426 gene partitions were further collapsed into 33 partitions by PartitionFinder v.2.1.1 with a greedy search to find partitions with consistent phylogenetic signals (Lanfear et al., 2012). Phylogenetic trees were constructed from this super-alignment and partition scheme with two methods—maximum likelihood implemented in IQ-TREE (v.1.5.5) and Bayesian inference implemented in BEAST (v.1.8.4) (Drummond and Rambaut, 2007; Nguyen et al., 2015). Configuration files for divergence time estimation analysis were coded in BEAUti v.1.8.4 using the 33 partitions and two calibration priors: 1) a direct fossil record of Chytridiomycota from the Rhynie Chert (407 Mya) (Krings et al., 2016; Strullu-Derrien et al., 2016), and 2) the emergence time of Chytridiomycota (573-770 Mya as 95% HPD) from earlier studies (Chang et al., 2015; Lutzoni et al., 2018; Wang et al., n.d.). The Birth-Death incomplete sampling tree model was employed for inter-species relationships analyses (Stadler, 2009). Unlinked strict clock models were used for each partition. Archive of input files and analysis scripts used to perform the phylogenetic analyses are available at Zenodo (DOI: 10.5281/zenodo.1447226). Three independent runs were performed separately for 50 million generations each with random starting seeds. Sufficient ESS (>200) values were obtained after the default burn-in (10%) for the final sampled trees. The Maximum Clade Credibility (MCC) tree was compiled using TreeAnnotator v.1.8.4.

### Identification of AGF-specific genes and Pfam domains

Orthologous genes across the 31 genomes or transcriptomes were identified using a comparative genomic pipeline that utilized all-vs-all BLASTp (cutoff=1E^-5^) to obtain the similarity pairs, Orthagogue to identify putative orthologous relationships, and the Markov-Clustering Algorithm (MCL using the inflation value of 1.5) to generate disjoint clusters and deployed in an analysis pipeline (DOI: 10.5281/zenodo.1447224) (Altschul et al., 1990; Ekseth et al., 2014; Van Dongen, 2000). Comparisons of shared gene content of the Orthologous clusters was performed among the Chytridiomycota lineages using a permissive strategy of counting a gene family as shared if it is missing in up to 5 of the 26 Neocallimastigomycota taxa and 1 of the 5 chytrids genomes. In this scenario, genes absent in all chytrids genomes and maintained by more than 21 out of the 26 Neocallimastigomycota genomes/transcriptomes are defined as AGF unique genes; on the other hand, genes missing from all Neocallimastigomycota and present in at least 4 out of the 5 chytrids genomes are treated as AGF lost genes.

Protein domains were identified by searching the predicted proteomes from each genome assembly or transcriptome assembly against the Protein Family (Pfam) database (v31.0, last accessed at March 20^th^, 2018). The enrichment heatmap of the Pfam domains across the included taxa was produced using the “aheatmap” function in the R package “NMF” based on the total copy number count in each assembly (Gaujoux and Seoighe, 2010). Genes only present in the AGF genomes and missing from all of the included free-living chytrids relatives were identified.

To identify genes in AGF that are likely important for interactions with mammalian hosts and plant material breakdown, we further compared the five available AGF genomes to the genomes of their animal hosts (e.g., sheep, horse, elephant, yak) (Broad Institute, 2018; Qiu et al., 2012; The International Sheep Genomics Consortium et al., 2010; Wade et al., 2009), the diet plant (e.g., moss, rice, palm, maize, sorghum) (Jiao et al., 2017; Martin et al., 2016; Paterson et al., 2009; Peng et al., 2013; Rensing et al., 2008; Singh et al., 2013; Swarbreck et al., 2008; The International Brachypodium Initiative et al., 2010; The Rice Annotation Project, 2007; Zimin et al., 2017) (Table S2), and the 1,165 available fungal genomes from the ongoing 1KFG project (Grigoriev et al., 2014; Spatafora, 2011; Spatafora et al., 2017; Stajich, 2017). To prioritize AGF genes that may have been laterally acquired from these hosts, a Python script (Wang et al., 2016) and similarity search tool BLAT (Kent, 2002) was applied to filter out genetic elements in AGF with higher similarity to animal or plant homologs than any fungal ones, excluding the AGF themselves. Candidate genes for lateral transfer were ranked by the combination of the two strategies. The candidate genes with an assigned functional or biological process annotation were analyzed with priority using phylogenetic reconstruction to infer their potential origin.

### Identification of homologous sequences and potential origin of HGT candidate loci

Three Pfam domains “Cthe_2159”, “Gal_Lectin”, and “Rhamnogal_lyase” were identified to be unique to the AGF genomes as compared to the Chytridiomycota fungi or all other fungal members. To predict the donor lineages for these putative HGT events, we searched more broadly for homologues in genome databases of Plant, Metazoa, Fungi, Bacteria, and Protists in EnsEMBL (v37) (Zerbino et al., 2018) via the web-implemented HMMER tool (https://www.ebi.ac.uk/Tools/hmmer/) (cutoff=1E^-3^). Additional fungal homologues were found by searching the DOE JGI’s MycoCosm database (Grigoriev et al., 2014; Spatafora, 2011). The profile Hidden Markov Model tool phmmer in the HMMer package (Eddy, 2011) was used to search for similar sequences in the 1,165 fungal genomes using the query of edge-trimmed domain sequences from *An. robustus* (cutoff=1E^-3^).

Members of the “RhgB_N” sequences were obtained from the Pfam database classified in the “RhgB_N” (PF09284) family (Finn et al., 2016) along with the N-terminal sequences of the rhamnogalacturonate lyase families A, B, and C from GenBank (Gomez-Cortecero et al., 2015; Hacquard et al., 2016; Yoshino-Yasuda et al., 2012). A single dataset of “RhgB_N” and “Rhamnogal_lyase” family members from animals, fungi, plants, and bacteria was constructed from these searches. Domain names were confirmed using NCBI’s conserved domain search tool (cutoff=1E^-5^) with unaligned FASTA sequences (Marchler-Bauer et al., 2017). Similarly, homologs of the “Gal_Lectin” and “Cthe_2159” were obtained by searching for similar sequences in the previously described genome databases and the categorized Pfam database (families of “Gal_Lectin (PF02140)” and “Cthe_2159 (PF14262)”). Homologous sequences containing the “Cthe_2159” domain were only identified in Archaea and Bacteria, while the AGF copies are the first eukaryotic representatives identified with this domain. Homologs of the flanking domain “Glyco_transf_34” was obtained similarly from EnsEMBL genome databases described above using the edge-trimmed domain sequence from *An. robustus* (cutoff=1E^-5^). Highly similar sequences (>90%) were filtered using CD-HIT v4.6.4 followed by multiple sequence alignment with MUSCLE v3.8.31 (Edgar, 2004; Fu et al., 2012).

### Phylogenetic trees of the HGT candidates

In total, 747 sequences of the rhamnogalacturanate degradation proteins (including both “Rhamnogal_lyase” and “RhgB_N”) were included in the alignment. For the other two domains, “Gal_Lectin” and “Cthe_2159”, the alignments include 297 and 234 unique variants respectively. The “Cthe_2159” domain containing genes in the 5 AGF genomes were aligned separately using MUSCLE v3.8.31 in Mesquite software (Edgar, 2004; Maddison and Maddison, 2007). Both the upstream and downstream flanking regions of the studied Pfam domain sequences were trimmed using the Mesquite software (Maddison and Maddison, 2007).

Selection of the appropriate substitutional model, the maximum-likelihood phylogenetic tree reconstruction, and the ultrafast bootstrapping (1000 replicates) were conducted using the IQ-TREE v1.5.5 package (Hoang et al., 2017; Kalyaanamoorthy et al., 2017; Nguyen et al., 2015).

## Supporting information

Combined supplementary materials

Supplementary results and associated figures

## Acknowledgements

This work was supported by National Science Foundation Grants (DEB-1557110 to J.E.S. and DEB-1557102 to N.Y. and M.E.). Y.W. acknowledges the Mycological Society of America for the Translational Mycology Postdoctoral Award. Data analyses were performed on the University of California Riverside High-Performance Computational Cluster supported by NSF DBI-1429826 and NIH S10-OD016290.

