## Supplementary material for "Molecular dating of the emergence of anaerobic rumen fungi and the impact of laterally acquired genes": Combined supplementary materials

*Short title*: Molecular dating and HGT of anaerobic gut fungi

*Authors*: Yan Wang[*,†], Noha Youssef[‡], M.B. Couger[§], Radwa Hanafy[‡], Mostafa Elshahed[‡], Jason E. Stajich[*,†]

*Affiliations*:

[*]Department of Microbiology and Plant Pathology, University of California, Riverside, Riverside, California, 92521 USA.

[†]Institute for Integrative Genome Biology, University of California, Riverside, Riverside, California, 92521 USA.

[‡]Department of Microbiology and Molecular Genetics, Oklahoma State University, Stillwater, Oklahoma, 74074 USA.

[§]High Performance Computing Center, Oklahoma State University, Stillwater, Oklahoma, 74074 USA.

*To whom correspondence may be addressed:*

Yan Wang, +1 951.386.5197,

Jason E. Stajich, +1 951.827.2363,

**This Supplementary Materials PDF file includes:**

Figures  S1-6

Tables S1-2

**Supplementary figure legend:**

**Fig. S1.** Maximum likelihood phylogenetic tree of Neocallimastigomycota using Chytridiomycota as the outgroup. All bootstrap values (out of 100) are labeled on the branches.

**Fig. S2.** Presence (dark gray) and absence (light gray) of the homologous gene families across the genomes (and transcriptomes) of Neocallimastigomycota and Chytridiomycota. The 4,824 gene families were selected as universal homologous genes that present at least 21 out of the 26 Neocallimastigomycota genomes (and transcriptomes) with missing no more than 1 of the 5 included Chytridiomycota genomes. In addition, it also includes the unique gene families that are strictly absence from all Chytridiomycota but encoded by the Neocallimastigomycota (missing no more than 5 out of the 26 taxa).

**Fig. S3.** Mid-point rooted phylogenetic tree of the "Cthe_2159" domain encoded by the Neocallimastigomycota (red). All 126 Neocallimastigomycota (AGF) copies form a single clade (red) indicating the HGT donor, Clostridiales bacterium C5EMF8 (an obligate rumen bacterium), with strong support of maximum likelihood bootstrap (98/100).

**Fig. S4.** Phylogenetic tree of the animal-like "Gal-Lectin" domain identified in Neocallimastigomycota. Clades are colored in consistence with the Figure 5 (Neocallimastigomycota in red, animals in blue, plants in green, and bacteria in brown).

**Fig. S5.** Phylogenetic tree of the AGF "Gal-Lectin" flanking domain "Glyco_transf_34" (rooted with bacterium, in purple). AGF homologs are colored in red clustering with other fungal taxa (in black). Plant homologs are in green and protists in brown.

**Fig. S6.** Mid-point rooted phylogenetic tree of the "Rhamnogal_lyase" domain encoded by the Neocallimastigomycota (red). Labels are consistent with the Figure 6.

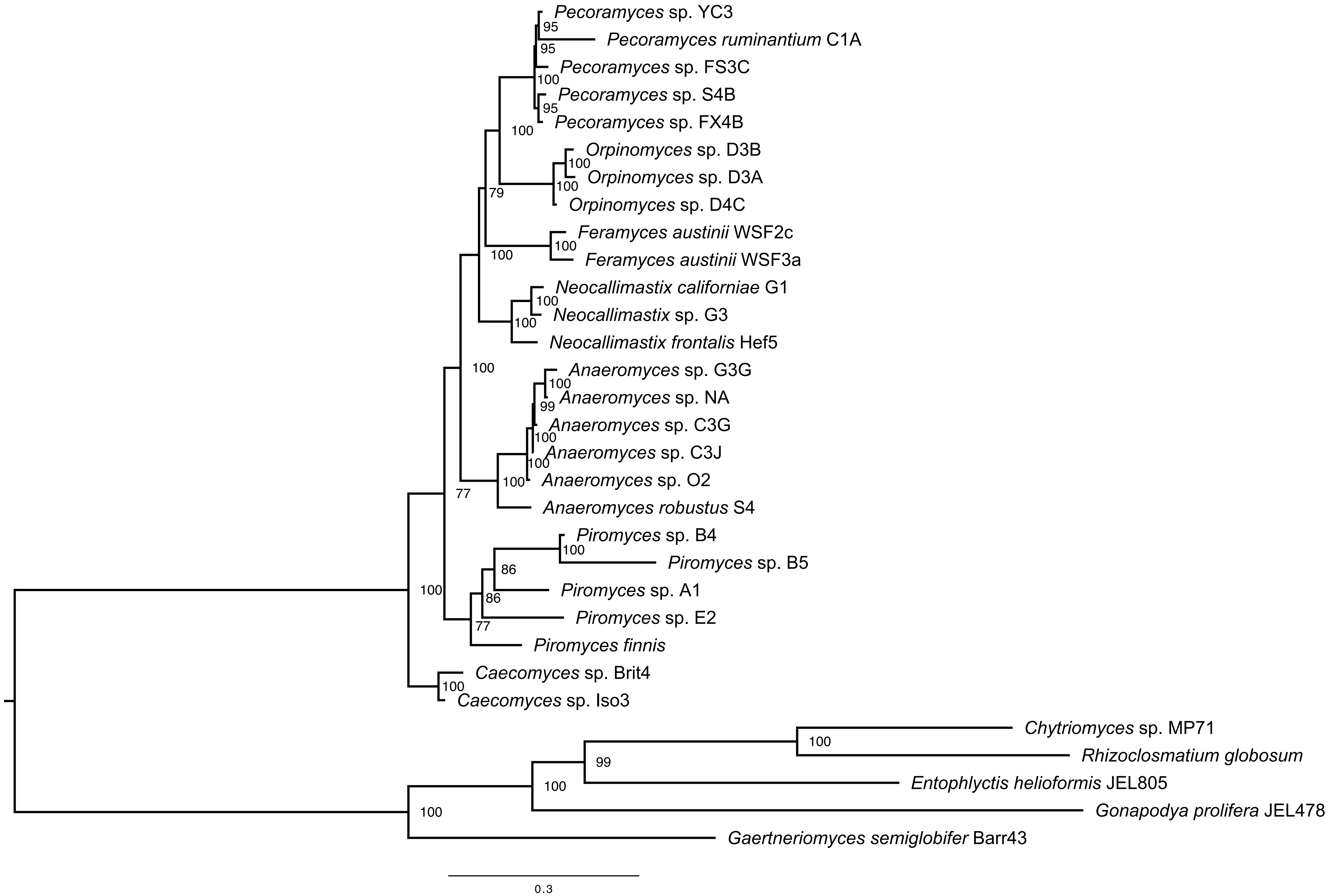
Pecoramyces sp. YC3
95
Pecoramyces ruminantium C1A
95
Pecoramyces sp. FS3C
100
Pecoramyces sp. S4B
95
Pecoramyces sp. FX4B
100
Orpinomyces sp. D3B
100
Orpinomyces sp. D3A
100
Orpinomyces sp. D4C
79
Feramyces austinii WSF2c
100
Feramyces austinii WSF3a
100
Neocallimastix californiae G1
100
Neocallimastix sp. G3
100
Neocallimastix frontalis Hef5
100
Anaeromyces sp. G3G
100
Anaeromyces sp. NA
99
Anaeromyces sp. C3G
100
Anaeromyces sp. C3J
100
Anaeromyces sp. O2
100
Anaeromyces robustus S4
77
Piromyces sp. B4
100
Piromyces sp. B5
86
Piromyces sp. A1
86
Piromyces sp. E2
77
Piromyces finnis
100
Caecomyces sp. Brit4
100
Caecomyces sp. Iso3
Chytriomyces sp. MP71
100
Rhizoclosmatium globosum
99
Entophlyctis helioformis JEL805
100
Gonapodya prolifera JEL478
100
Gaertneriomyces semiglobifer Barr43
0.3

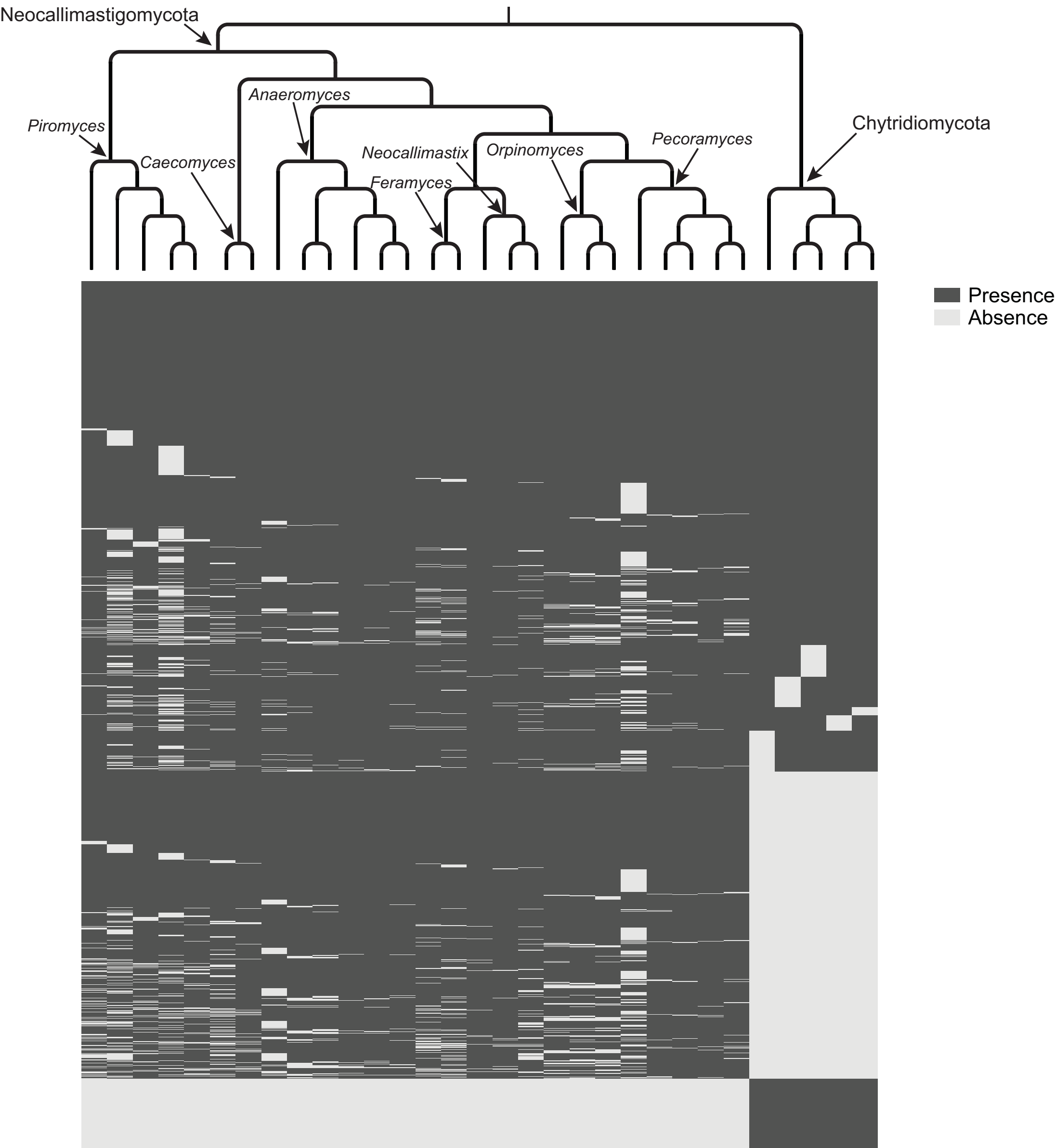

Neocallimastigomycota
Piromyces
Caecomyces
Anaeromyces
Neocallimastix
Orpinomyces
Pecoramyces
Feramyces
Chytridiomycota
Presence
Absence

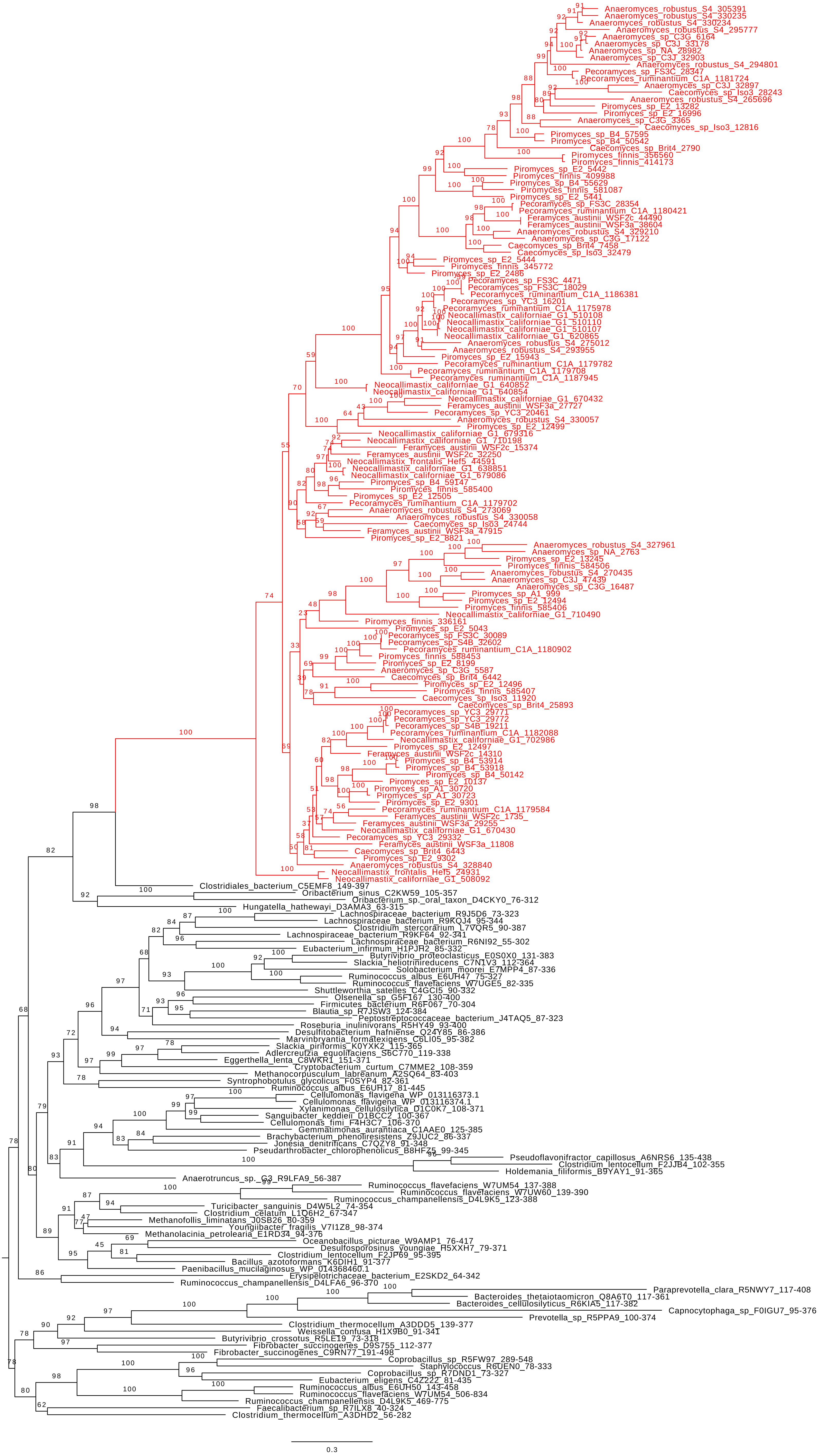
Anaeromyces_robustus_S4_305391
Anaeromyces_robustus_S4_330235
Anaeromyces_robustus_S4_330234
Anaeromyces_robustus_S4_295777
Anaeromyces_sp_C3G_6164
Anaeromyces_sp_C3J_33178
Anaeromyces_sp_NA_28982
Anaeromyces_sp_C3J_32903
Anaeromyces_robustus_S4_294801
Pecoramyces_sp_FS3C_28347
Pecoramyces_ruminantium_C1A_1181724
Anaeromyces_sp_C3J_32897
Caecomyces_sp_Iso3_28243
Anaeromyces_robustus_S4_265696
Piromyces_sp_E2_13282
Piromyces_sp_E2_16996
Anaeromyces_sp_C3G_3365
Caecomyces_sp_Iso3_12816
Piromyces_sp_B4_57595
Piromyces_sp_B4_50542
Caecomyces_sp_Brit4_2790
Piromyces_finnis_356560
Piromyces_finnis_414173
Piromyces_sp_E2_5442
Piromyces_finnis_409988
Piromyces_sp_B4_55629
Piromyces_finnis_581087
Piromyces_sp_E2_5441
Pecoramyces_sp_FS3C_28354
Pecoramyces_ruminantium_C1A_1180421
Feramyces_austinii_WSF2c_44490
Feramyces_austinii_WSF3a_38604
Anaeromyces_robustus_S4_329210
Anaeromyces_sp_C3G_17122
Caecomyces_sp_Brit4_7458
Caecomyces_sp_Iso3_32479
Piromyces_sp_E2_5444
Piromyces_finnis_345772
Piromyces_sp_E2_2486
Pecoramyces_sp_FS3C_4471
Pecoramyces_sp_FS3C_18029
Pecoramyces_ruminantium_C1A_1186381
Pecoramyces_sp_YC3_16201
Pecoramyces_ruminantium_C1A_1175978
Neocallimastix_californiae_G1_510108
Neocallimastix_californiae_G1_510110
Neocallimastix_californiae_G1_510107
Neocallimastix_californiae_G1_620865
Anaeromyces_robustus_S4_275012
Anaeromyces_robustus_S4_293955
Piromyces_sp_E2_15943
Pecoramyces_ruminantium_C1A_1179782
Pecoramyces_ruminantium_C1A_1179708
Pecoramyces_ruminantium_C1A_1187945
Neocallimastix_californiae_G1_640852
Neocallimastix_californiae_G1_640854
Neocallimastix_californiae_G1_670432
Feramyces_austinii_WSF3a_27727
Pecoramyces_sp_YC3_20461
Anaeromyces_robustus_S4_330057
Piromyces_sp_E2_12499
Neocallimastix_californiae_G1_679316
Neocallimastix_californiae_G1_710198
Feramyces_austinii_WSF2c_15374
Feramyces_austinii_WSF2c_32250
Neocallimastix_frontalis_Hef5_44591
Neocallimastix_californiae_G1_638851
Neocallimastix_californiae_G1_679086
Piromyces_sp_B4_59147
Piromyces_finnis_585400
Piromyces_sp_E2_12505
Pecoramyces_ruminantium_C1A_1179702
Anaeromyces_robustus_S4_273069
Anaeromyces_robustus_S4_330058
Caecomyces_sp_Iso3_24744
Feramyces_austinii_WSF3a_47915
Piromyces_sp_E2_8821
Anaeromyces_robustus_S4_327961
Anaeromyces_sp_NA_2763
Piromyces_sp_E2_13245
Piromyces_finnis_584506
Anaeromyces_robustus_S4_270435
Anaeromyces_sp_C3J_47439
Anaeromyces_sp_C3G_16487
Piromyces_sp_A1_999
Piromyces_sp_E2_12494
Piromyces_finnis_585406
Neocallimastix_californiae_G1_710490
Piromyces_finnis_336161
Piromyces_sp_E2_5043
Pecoramyces_sp_FS3C_30089
Pecoramyces_sp_S4B_32602
Pecoramyces_ruminantium_C1A_1180902
Piromyces_finnis_588453
Piromyces_sp_E2_8199
Anaeromyces_sp_C3G_5587
Caecomyces_sp_Brit4_6442
Piromyces_sp_E2_12496
Piromyces_finnis_585407
Caecomyces_sp_Iso3_11920
Caecomyces_sp_Brit4_25893
Pecoramyces_sp_YC3_29771
Pecoramyces_sp_YC3_29772
Pecoramyces_sp_S4B_19211
Pecoramyces_ruminantium_C1A_1182088
Neocallimastix_californiae_G1_702986
Piromyces_sp_E2_12497
Feramyces_austinii_WSF2c_14310
Piromyces_sp_B4_53914
Piromyces_sp_B4_53918
Piromyces_sp_B4_50142
Piromyces_sp_E2_10137
Piromyces_sp_A1_30720
Piromyces_sp_A1_30723
Piromyces_sp_E2_9301
Pecoramyces_ruminantium_C1A_1179584
Feramyces_austinii_WSF2c_1735_
Feramyces_austinii_WSF3a_29255
Neocallimastix_californiae_G1_670430
Pecoramyces_sp_YC3_29332
Feramyces_austinii_WSF3a_11808
Caecomyces_sp_Brit4_6443
Piromyces_sp_E2_9302
Anaeromyces_robustus_S4_328840
Neocallimastix_frontalis_Hef5_24931
Neocallimastix_californiae_G1_508092
Clostridiales_bacterium_C5EMF8_149-397
Oribacterium_sinus_C2KW59_105-357
Oribacterium_sp._oral_taxon_D4CKY0_76-312
Hungatella_hathewayi_D3AMA3_63-315
Lachnospiraceae_bacterium_R9J5D6_73-323
Lachnospiraceae_bacterium_R9KQJ4_95-344
Clostridium_stercorarium_L7VQR5_90-387
Lachnospiraceae_bacterium_R9KF64_92-341
Lachnospiraceae_bacterium_R6NI92_55-302
Eubacterium_infirmum_H1PJH2_85-332
Butyrivibrio_proteoclasticus_E0S0X0_131-383
Slackia_heliotrinireducens_C7N1V3_112-364
Solobacterium_moorei_E7MPP4_87-336
Ruminococcus_albus_E6UH47_75-327
Ruminococcus_flavefaciens_W7UGE5_82-335
Shuttleworthia_satelles_C4GCI5_90-332
Olsenella_sp_G5F167_130-400
Firmicutes_bacterium_R6F067_70-304
Blautia_sp_R7JSW3_124-384
Peptostreptococcaceae_bacterium_J4TAQ5_87-323
Roseburia_inulinivorans_R5HY49_93-400
Desulfitobacterium_hafniense_Q24Y85_86-386
Marvinbryantia_formatexigens_C6LI05_95-382
Slackia_piriformis_K0YXK2_115-365
Adlercreutzia_equolifaciens_S6C770_119-338
Eggerthella_lenta_C8WKR1_151-371
Cryptobacterium_curtum_C7MME2_108-359
Methanocorpusculum_labreanum_A2SQ64_83-403
Syntrophobotulus_glycolicus_F0SYP4_82-361
Ruminococcus_albus_E6UR17_81-445
Cellulomonas_flavigena_WP_013116373.1
Cellulomonas_flavigena_WP_013116374.1
Xylanimonas_cellulosilytica_D1C0K7_108-371
Sanguibacter_keddieii_D1BCC2_100-367
Cellulomonas_fimi_F4H3C7_106-370
Gemmatimonas_aurantiaca_C1AAE0_125-385
Brachybacterium_phenoliresistens_Z9JUC2_86-337
Jonesia_denitrificans_C7QZY8_91-348
Pseudarthrobacter_chlorophenolicus_B8HFZ5_99-345
Pseudoflavonifractor_capillosus_A6NRS6_135-438
Clostridium_lentocellum_F2JJB4_102-355
Holdemania_filiformis_B9YAY1_91-365
Anaerotruncus_sp._G3_R9LFA9_56-387
Ruminococcus_flavefaciens_W7UM54_137-388
Ruminococcus_flavefaciens_W7UW60_139-390
Ruminococcus_champanellensis_D4L9K5_123-388
Turicibacter_sanguinis_D4W5L2_74-354
Clostridium_celatum_L1Q6H2_67-347
Methanofollis_liminatans_J0SB26_80-359
Youngiibacter_fragilis_V7I1Z8_98-374
Methanolacinia_petrolearia_E1RD34_94-376
Oceanobacillus_picturae_W9AMP1_76-417
Desulfosporosinus_youngiae_H5XXH7_79-371
Clostridium_lentocellum_F2JP69_95-395
Bacillus_azotoformans_K6DIH1_91-377
Paenibacillus_mucilaginosus_WP_014368460.1
Erysipelotrichaceae_bacterium_E2SKD2_64-342
Ruminococcus_champanellensis_D4LFA6_96-370
Paraprevotella_clara_R5NWY7_117-408
Bacteroides_thetaiotaomicron_Q8A6T0_117-361
Bacteroides_cellulosilyticus_R6KIA5_117-382
Capnocytophaga_sp_F0IGU7_95-376
Prevotella_sp_R5PPA9_100-374
Clostridium_thermocellum_A3DDD5_139-377
Weissella_confusa_H1X9B0_91-341
Butyrivibrio_crossotus_R5LE19_73-318
Fibrobacter_succinogenes_D9S755_112-377
Fibrobacter_succinogenes_C9RN77_191-498
Coprobacillus_sp_R5FW97_289-548
Staphylococcus_R6UEN0_78-333
Coprobacillus_sp_R7DND1_73-327
Eubacterium_eligens_C4Z222_81-435
Ruminococcus_albus_E6UH50_143-458
Ruminococcus_flavefaciens_W7UM54_506-834
Ruminococcus_champanellensis_D4L9K5_469-775
Faecalibacterium_sp_R7ILX8_40-324
Clostridium_thermocellum_A3DHD2_56-282
0.3

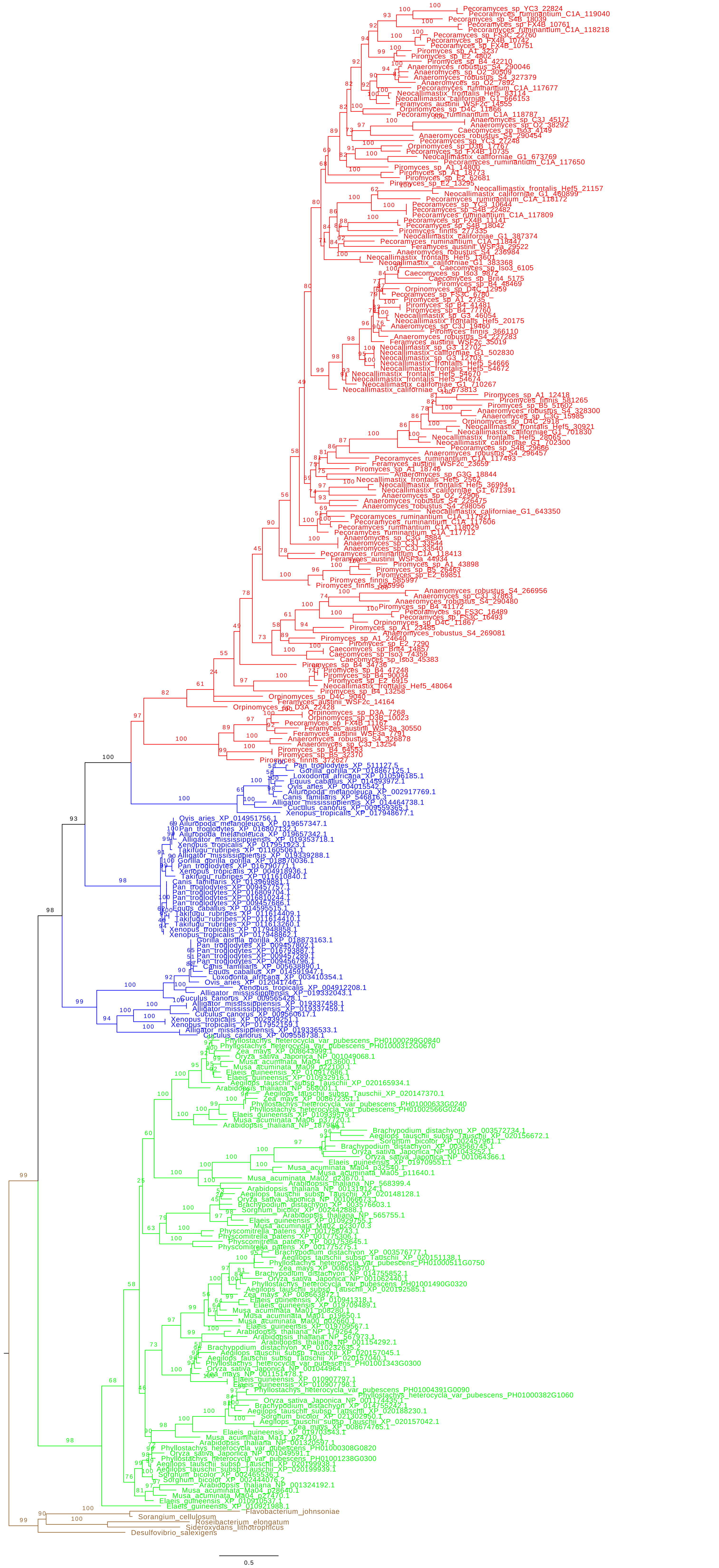
Flavobacterium_johnsoniae
Sorangium_cellulosum
Roseibacterium_elongatum
Sideroxydans_lithotrophicus
Desulfovibrio_salexigens
0.5

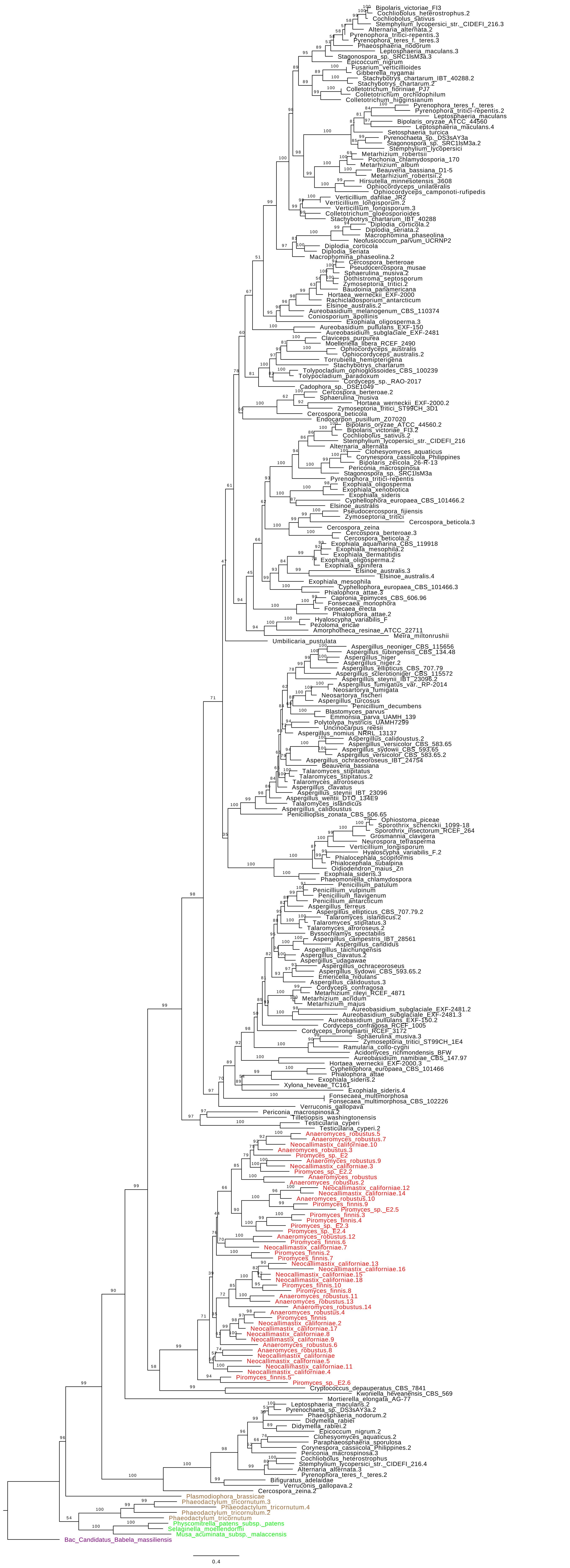
Bipolaris_victoriae_FI3
Cochliobolus_heterostrophus.2
Cochliobolus_sativus
Stemphylium_lycopersici_str._CIDEFI_216.3
Alternaria_alternata.2
Pyrenophora_tritici-repentis.3
Pyrenophora_teres_f._teres.3
Phaeosphaeria_nodorum
Leptosphaeria_maculans.3
Stagonospora_sp._SRC1lsM3a.3
Epicoccum_nigrum
Fusarium_verticillioides
Gibberella_nygamai
Stachybotrys_chartarum_IBT_40288.2
Stachybotrys_chartarum.2
Colletotrichum_fioriniae_PJ7
Colletotrichum_orchidophilum
Colletotrichum_higginsianum
Pyrenophora_teres_f._teres
Pyrenophora_tritici-repentis.2
Leptosphaeria_maculans
Bipolaris_oryzae_ATCC_44560
Leptosphaeria_maculans.4
Setosphaeria_turcica
Pyrenochaeta_sp._DS3sAY3a
Stagonospora_sp._SRC1lsM3a.2
Stemphylium_lycopersici
Metarhizium_robertsii
Pochonia_chlamydosporia_170
Metarhizium_album
Beauveria_bassiana_D1-5
Metarhizium_robertsii.2
Hirsutella_minnesotensis_3608
Ophiocordyceps_unilateralis
Ophiocordyceps_camponoti-rufipedis
Verticillium_dahliae_JR2
Verticillium_longisporum.2
Verticillium_longisporum.3
Colletotrichum_gloeosporioides
Stachybotrys_chartarum_IBT_40288
Diplodia_corticola.2
Diplodia_seriata.2
Macrophomina_phaseolina
Neofusicoccum_parvum_UCRNP2
Diplodia_corticola
Diplodia_seriata
Macrophomina_phaseolina.2
Cercospora_berteroae
Pseudocercospora_musae
Sphaerulina_musiva.2
Dothistroma_septosporum
Zymoseptoria_tritici.2
Baudoinia_panamericana
Hortaea_werneckii_EXF-2000
Rachicladosporium_antarcticum
Elsinoe_australis.2
Aureobasidium_melanogenum_CBS_110374
Coniosporium_apollinis
Exophiala_oligosperma.3
Aureobasidium_pullulans_EXF-150
Aureobasidium_subglaciale_EXF-2481
Claviceps_purpurea
Moelleriella_libera_RCEF_2490
Ophiocordyceps_australis
Ophiocordyceps_australis.2
Torrubiella_hemipterigena
Stachybotrys_chartarum
Tolypocladium_ophioglossoides_CBS_100239
Tolypocladium_paradoxum
Cordyceps_sp._RAO-2017
Cadophora_sp._DSE1049
Cercospora_berteroae.2
Sphaerulina_musiva
Hortaea_werneckii_EXF-2000.2
Zymoseptoria_tritici_ST99CH_3D1
Cercospora_beticola
Endocarpon_pusillum_Z07020
Bipolaris_oryzae_ATCC_44560.2
Bipolaris_victoriae_FI3.2
Cochliobolus_sativus.2
Stemphylium_lycopersici_str._CIDEFI_216
Alternaria_alternata
Clohesyomyces_aquaticus
Corynespora_cassiicola_Philippines
Bipolaris_zeicola_26-R-13
Periconia_macrospinosa
Stagonospora_sp._SRC1lsM3a
Pyrenophora_tritici-repentis
Exophiala_oligosperma
Exophiala_xenobiotica
Exophiala_sideris
Cyphellophora_europaea_CBS_101466.2
Elsinoe_australis
Pseudocercospora_fijiensis
Zymoseptoria_tritici
Cercospora_beticola.3
Cercospora_zeina
Cercospora_berteroae.3
Cercospora_beticola.2
Exophiala_aquamarina_CBS_119918
Exophiala_mesophila.2
Exophiala_dermatitidis
Exophiala_oligosperma.2
Exophiala_spinifera
Elsinoe_australis.3
Elsinoe_australis.4
Exophiala_mesophila
Cyphellophora_europaea_CBS_101466.3
Phialophora_attae.3
Capronia_epimyces_CBS_606.96
Fonsecaea_monophora
Fonsecaea_erecta
Phialophora_attae.2
Hyaloscypha_variabilis_F
Pezoloma_ericae
Amorphotheca_resinae_ATCC_22711
Meira_miltonrushii
Umbilicaria_pustulata
Aspergillus_neoniger_CBS_115656
Aspergillus_tubingensis_CBS_134.48
Aspergillus_niger
Aspergillus_niger.2
Aspergillus_ellipticus_CBS_707.79
Aspergillus_sclerotioniger_CBS_115572
Aspergillus_steynii_IBT_23096.2
Aspergillus_fumigatus_var._RP-2014
Neosartorya_fumigata
Neosartorya_fischeri
Aspergillus_turcosus
Penicillium_decumbens
Blastomyces_parvus
Emmonsia_parva_UAMH_139
Polytolypa_hystricis_UAMH7299
Uncinocarpus_reesii
Aspergillus_nomius_NRRL_13137
Aspergillus_calidoustus.2
Aspergillus_versicolor_CBS_583.65
Aspergillus_sydowii_CBS_593.65
Aspergillus_versicolor_CBS_583.65.2
Aspergillus_ochraceoroseus_IBT_24754
Beauveria_bassiana
Talaromyces_stipitatus
Talaromyces_stipitatus.2
Talaromyces_atroroseus
Aspergillus_clavatus
Aspergillus_steynii_IBT_23096
Aspergillus_wentii_DTO_134E9
Talaromyces_islandicus
Aspergillus_calidoustus
Penicilliopsis_zonata_CBS_506.65
Ophiostoma_piceae
Sporothrix_schenckii_1099-18
Sporothrix_insectorum_RCEF_264
Grosmannia_clavigera
Neurospora_tetrasperma
Verticillium_longisporum
Hyaloscypha_variabilis_F.2
Phialocephala_scopiformis
Phialocephala_subalpina
Oidiodendron_maius_Zn
Exophiala_sideris.3
Phaeomoniella_chlamydospora
Penicillium_patulum
Penicillium_vulpinum
Penicillium_flavigenum
Penicillium_antarcticum
Aspergillus_terreus
Aspergillus_ellipticus_CBS_707.79.2
Talaromyces_islandicus.2
Talaromyces_stipitatus.3
Talaromyces_atroroseus.2
Byssochlamys_spectabilis
Aspergillus_campestris_IBT_28561
Aspergillus_candidus
Aspergillus_taichungensis
Aspergillus_clavatus.2
Aspergillus_udagawae
Aspergillus_ochraceoroseus
Aspergillus_sydowii_CBS_593.65.2
Emericella_nidulans
Aspergillus_calidoustus.3
Cordyceps_confragosa
Metarhizium_rileyi_RCEF_4871
Metarhizium_acridum
Metarhizium_majus
Aureobasidium_subglaciale_EXF-2481.2
Aureobasidium_subglaciale_EXF-2481.3
Aureobasidium_pullulans_EXF-150.2
Cordyceps_confragosa_RCEF_1005
Cordyceps_brongniartii_RCEF_3172
Sphaerulina_musiva.3
Zymoseptoria_tritici_ST99CH_1E4
Ramularia_collo-cygni
Acidomyces_richmondensis_BFW
Aureobasidium_namibiae_CBS_147.97
Hortaea_werneckii_EXF-2000.3
Cyphellophora_europaea_CBS_101466
Phialophora_attae
Exophiala_sideris.2
Xylona_heveae_TC161
Exophiala_sideris.4
Fonsecaea_multimorphosa
Fonsecaea_multimorphosa_CBS_102226
Verruconis_gallopava
Periconia_macrospinosa.2
Tilletiopsis_washingtonensis
Testicularia_cyperi
Testicularia_cyperi.2
Anaeromyces_robustus.5
Anaeromyces_robustus.7
Neocallimastix_californiae.10
Anaeromyces_robustus.3
Piromyces_sp._E2
Anaeromyces_robustus.9
Neocallimastix_californiae.3
Piromyces_sp._E2.2
Anaeromyces_robustus
Anaeromyces_robustus.2
Neocallimastix_californiae.12
Neocallimastix_californiae.14
Anaeromyces_robustus.10
Piromyces_finnis.9
Piromyces_sp._E2.5
Piromyces_finnis.3
Piromyces_finnis.4
Piromyces_sp._E2.3
Piromyces_sp._E2.4
Anaeromyces_robustus.12
Piromyces_finnis.6
Neocallimastix_californiae.7
Piromyces_finnis.2
Piromyces_finnis.7
Neocallimastix_californiae.13
Neocallimastix_californiae.16
Neocallimastix_californiae.15
Neocallimastix_californiae.18
Piromyces_finnis.10
Piromyces_finnis.8
Anaeromyces_robustus.11
Anaeromyces_robustus.13
Anaeromyces_robustus.14
Anaeromyces_robustus.4
Piromyces_finnis
Neocallimastix_californiae.2
Neocallimastix_californiae.17
Neocallimastix_californiae.8
Neocallimastix_californiae.9
Anaeromyces_robustus.6
Anaeromyces_robustus.8
Neocallimastix_californiae
Neocallimastix_californiae.5
Neocallimastix_californiae.11
Neocallimastix_californiae.4
Piromyces_finnis.5
Piromyces_sp._E2.6
Cryptococcus_depauperatus_CBS_7841
Kwoniella_heveanensis_CBS_569
Mortierella_elongata_AG-77
Leptosphaeria_maculans.2
Pyrenochaeta_sp._DS3sAY3a.2
Phaeosphaeria_nodorum.2
Didymella_rabiei
Didymella_rabiei.2
Epicoccum_nigrum.2
Clohesyomyces_aquaticus.2
Paraphaeosphaeria_sporulosa
Corynespora_cassiicola_Philippines.2
Periconia_macrospinosa.3
Cochliobolus_heterostrophus
Stemphylium_lycopersici_str._CIDEFI_216.4
Alternaria_alternata.3
Pyrenophora_teres_f._teres.2
Bifiguratus_adelaidae
Verruconis_gallopava.2
Cercospora_zeina.2
Plasmodiophora_brassicae
Phaeodactylum_tricornutum.3
Phaeodactylum_tricornutum.4
Phaeodactylum_tricornutum.2
Phaeodactylum_tricornutum
Physcomitrella_patens_subsp._patens
Selaginella_moellendorffii
Musa_acuminata_subsp._malaccensis
Bac_Candidatus_Babela_massiliensis
0.4

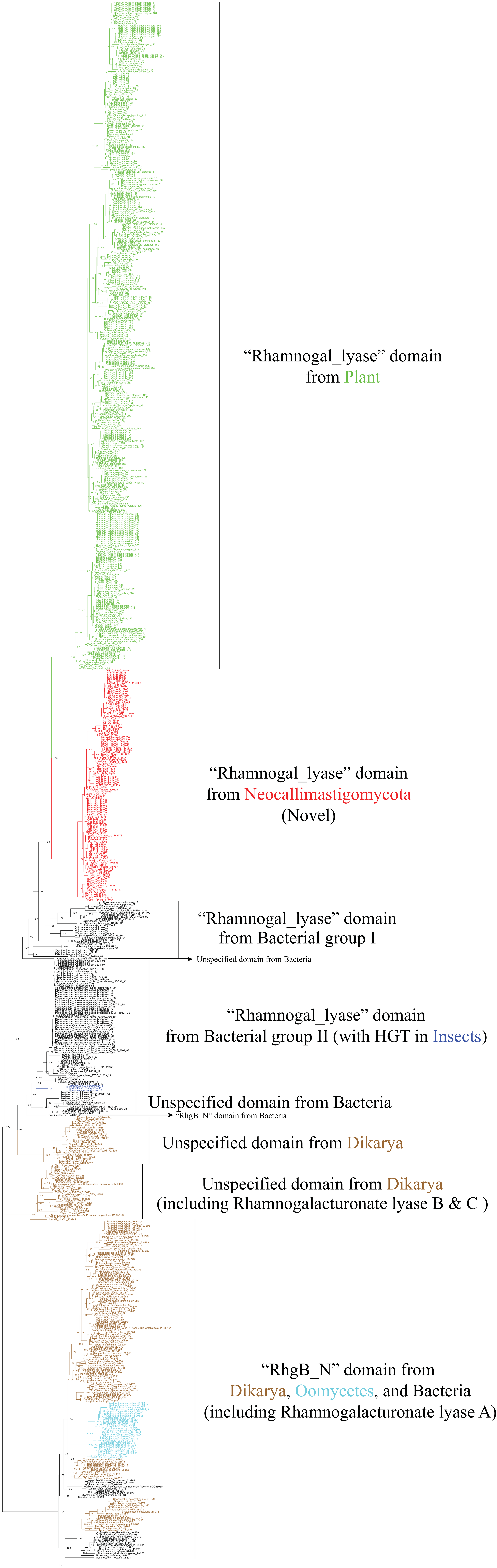

“Rhamnogal_lyase” domain
from Plant
“Rhamnogal_lyase” domain
from Neocallimastigomycota
(Novel)
“Rhamnogal_lyase” domain
from Bacterial group I
Unspecified domain from Bacteria
“Rhamnogal_lyase” domain
from Bacterial group II (with HGT in Insects)
Unspecified domain from Bacteria
“RhgB_N” domain from Bacteria
Unspecified domain from Dikarya
Unspecified domain from Dikarya
(including Rhamnogalacturonate lyase B & C )
“RhgB_N” domain from
Dikarya, Oomycetes, and Bacteria
(including Rhamnogalacturonate lyase A)

**Table S1.** List of Pfam domain names with annotated functions for the ones uniquely maintained or lost in Neocallimastigomycota (AGF) comparing to Chytridiomycota

| **AGF Unique Domains** | | **AGF Lost Domains** | |
|---|---|---|---|
| Domain | Function | Domain | Function |
| AAT | Acyl-coenzyme A:6-aminopenicillanic acid acyl-transferase | 2OG-FeII_Oxy_3 | 2OG-Fe(II) oxygenase superfamily |
| Abhydrolase_5 | Alpha/beta hydrolase superfamily, hydrolytic enzymes of widely differing phylogenetic origin and catalytic function that share a common fold. | 2OG-FeII_Oxy_4 | 2OG-Fe(II) oxygenase superfamily |
| Asp_protease | Aspartic proteases are a catalytic type of protease enzymes that use an activated water molecule bound to one or more aspartate residues for catalysis of their peptide substrates. | 3-HAO | 3-hydroxyanthranilic acid dioxygenase, part of the kynurenine pathway for the degradation of tryptophan and the biosynthesis of nicotinic acid |
| Asp_protease_2 | Aspartic proteases | ACAS_N | Acetyl-coenzyme A synthetase N-terminus |
| Aspzincin_M35 | This is the catalytic region of aspzincins, a group of lysine-specific metallo-endopeptidases in the M35 family | Alg6_Alg8 | N-linked (asparagine-linked) glycosylation of proteins is mediated by a highly conserved pathway in eukaryotes, in which a lipid (dolichol phosphate)-linked oligosaccharide is assembled at the endoplasmic reticulum membrane prior to the transfer of the oligosaccharide moiety to the target asparagine residues. |
| Carb_bind | This is a carbohydrate binding domain which has been shown in Schizosaccharomyces pombe to be required for septum localisation | Allantoicase | This enzyme belongs to the family of hydrolases, those acting on carbon-nitrogen bonds other than peptide bonds, specifically in linear amidines. The systematic name of this enzyme class is allantoate amidinohydrolase. This enzyme participates in purine metabolism by facilitating the utilization of purines as secondary nitrogen sources under nitrogen-limiting conditions. While purine degradation converges to uric acid in all vertebrates, its further degradation varies from species to species. Amphibians and microorganisms produce ammonia and carbon dioxide using the uricolytic pathway. Allantoicase performs the second step in this pathway catalyzing the conversion of allantoate into ureidoglycolate and urea. |
| CBAH | Choloylglycine hydrolase family | ATP_sub_h | ATP synthase complex subunit h |
| CBM_10 | Carbohydrate-binding module family 10 (CBM10) is found in two distinct sets of proteins with different functions. Those found in aerobic bacteria bind cellulose (or other carbohydrates); but in anaerobic fungi they are protein binding | ATP-synt_10 | mitochondrial ATPase complex subunit ATP1 |

| | | | |
|---|---|---|---|
| | domains, referred to as dockerin domains. The dockerin domains are believed to be responsible for the assembly of a multiprotein cellulase/hemicellulase complex, similar to the cellulosome found in certain anaerobic bacteria. | | |
| CBM_25 | Carbohydrate-binding module family 25 (CBM25) binds alpha-glucooligosaccharides, particularly those containing alpha-1,6 linkages, and granular starch. | ATP11 | Atp11p is a molecular chaperone of the mitochondrial matrix that participates in the biogenesis pathway to form F1, the catalytic unit of the ATP synthase |
| CBM_35 | This is a mannan-specific carbohydrate binding domain, previously known as the X4 module. Unlike other carbohydrate binding modules, binding to substrate causes a conformational change | B12D | NADH dehydrogenase (ubiquinone) |
| CBM_6 | Carbohydrate-binding module family 6 (CBM6) is unusual in that is contains two substrate-binding sites, cleft A and cleft B. Cellvibrio mixtus endoglucanase 5A contains two CBM6 domains, the CBM6 domain at the C-terminus displays distinct ligand binding specificities in each of the sustrate-binding clefts. Both cleft A and cleft B can bind cello-oligosaccharides, laminarin preferentially binds in cleft A, xylooligosaccharides only bind in cleft A and beta1,4,-beta1,3-mixed linked glucans only bind in cleft B. | Band_7_C | C-terminal region of band_7 |
| CBM-like | Polysaccharide lyase family 4, domain III | BCDHK_Adom3 | Mitochondrial branched-chain alpha-ketoacid dehydrogenase kinase |
| CBM26 | CBM26 is a carbohydrate-binding module that binds starch | BCS1_N | Mitochondrial chaperone BCS1 |
| CHB_HEX_C_1 | Chitobiase/beta-hexosaminidase C-terminal domain | Bot1p | Eukaryotic mitochondrial regulator protein |
| CotH | CotH kinase protein, members of this family include the spore coat protein H (cotH). This protein is an atypical protein kinase that phosphorylates CotB and CotG | Chalcone_2 | Chalcone isomerase like |
| Cthe_2159 | Carbohydrate-binding domain-containing protein | Chromate_transp | Members of this family probably act as chromate transporters. Members of this family are found in both bacteria and archaebacteria. The proteins are composed of one or two copies of this region. The alignment contains two conserved motifs, FGG and PGP. |
| FeoA | This family includes FeoA a small protein, probably involved in Fe2+ transport [1]. This presumed short domain is also found at the C-terminus of a variety of metal dependent | CIAPIN1 | Cytokine-induced anti-apoptosis inhibitor 1, Fe-S biogenesis. Anamorsin, subsequently named CIAPIN1 for cytokine-induced anti-apoptosis inhibitor 1, in humans is the homologue of yeast |

| | | | |
|---|---|---|---|
| | transcriptional regulators. This suggests that this domain may be metal-binding. In most cases this is likely to be either iron or manganese. | | Dre2, a conserved soluble eukaryotic Fe-S cluster protein, that functions in cytosolic Fe-S protein biogenesis. It is found in both the cytoplasm and in the mitochondrial intermembrane space (IMS) |
| FeoB_C | Ferrous iron transport protein B C terminus | Cmc1 | Cytochrome c oxidase biogenesis protein Cmc1 like |
| FLYWCH | In molecular biology, the FLYWCH zinc finger is a zinc finger domain. It is found in a number of eukaryotic proteins. FLYWCH is a C2H2-type zinc finger characterised by five conserved hydrophobic residues, containing the conserved sequence motif: | Coa1 | Cytochrome oxidase complex assembly protein 1 |
| Gal_Lectin | Galactose binding lectin domain | Complex1_LYR | NADH dehydrogenase (ubiquinone) |
| Glyco_hydro_11 | Glycoside hydrolase family 11 CAZY GH_11 comprises enzymes with only one known activity, xylanase (EC 3.2.1.8). These enzymes were formerly known as cellulase family G | Complex1_LYR_2 | This is a family of proteins carrying the LYR motif of family Complex1_LYR, PF05347 likely to be involved in Fe-S cluster biogenesis in mitochondria. |
| Glyco_hydro_48 | Glycoside hydrolase family 48 CAZY GH_48 comprises enzymes with several known activities; endoglucanase (EC 3.2.1.4); cellobiohydrolase (EC 3.2.1.91). | COQ7 | Ubiquinone biosynthesis protein COQ7 |
| Glyco_hydro_6 | Glycoside hydrolase family 6 comprises enzymes with several known activities including endoglucanase (EC 3.2.1.4) and cellobiohydrolase (EC 3.2.1.91). These enzymes were formerly known as cellulase family B. The 3D structure of the enzymatic core of cellobiohydrolase II (CBHII) from the fungus Trichoderma reesei reveals an alpha-beta protein with a fold similar to the ubiquitous barrel topology first seen in triose phosphate isomerase. | COX15-CtaA | Cytochrome c oxidase |
| Glyco_hydro_8 | Glycoside hydrolase family 8 CAZY GH_8 comprises enzymes with several known activities; endoglucanase (EC 3.2.1.4); lichenase (EC 3.2.1.73); chitosanase (EC 3.2.1.132). These enzymes were formerly known as cellulase family D. | COX6B | Cytochrome c oxidase, the last enzyme in the respiratory electron transport chain of mitochondria (or bacteria) located in the mitochondrial (or bacterial) membrane. |
| GT-D | Glycosyltransferase GT-D fold | CtaG_Cox11 | Cytochrome c oxidase |
| LRR_5 | A leucine-rich repeat (LRR) is a protein structural motif that forms an α/β horseshoe fold. It is composed of repeating 20–30 amino acid stretches that are unusually rich in the hydrophobic amino acid leucine. These repeats commonly fold together to form | Cupin_4 | Cupin superfamily protein |

| | | | |
|---|---|---|---|
| | a solenoid protein domain, termed leucine-rich repeat domain. Typically, each repeat unit has beta strand-turn-alpha helix structure, and the assembled domain, composed of many such repeats, has a horseshoe shape with an interior parallel beta sheet and an exterior array of helices. One face of the beta sheet and one side of the helix array are exposed to solvent and are therefore dominated by hydrophilic residues. The region between the helices and sheets is the protein's hydrophobic core and is tightly sterically packed with leucine residues.<br>Leucine-rich repeats are frequently involved in the formation of protein–protein interactions. | | |
| NRDD | Anaerobic ribonucleoside-triphosphate reductase | Cupin_8 | This cupin like domain shares similarity to the JmjC domain, which catalyse a novel histone modification |
| PDDEXK_2 | PD-(D/E)XK nuclease family transposase. These proteins are transposase proteins. | Cyto_heme_lyase | Holocytochrome-c synthase |
| Pollen_allerg_1 | This family contains allergens lol PI, PII and PIII from Lolium perenne. | Cytochrom_B561 | Cytochrome b561 is an integral membrane protein responsible for electron transport, binding two heme groups non-covalently.[1] It is a family of ascorbate-dependent oxidoreductase enzymes |
| Pur_ac_phosph_N | Purple acid Phosphatase, N-terminal domain | DAP3 | Mitochondrial ribosomal death-associated protein 3 |
| Recombinase | This domain is usually found associated with PF00239 in putative integrases/recombinases of mobile genetic elements of diverse bacteria and phages. | DNA_photolyase | Deoxyribodipyrimidine photolyase (DNA photolyase) is a DNA repair enzyme. It binds to UV-damaged DNA containing pyrimidine dimers and, upon absorbing a near-UV photon (300 to 500 nm), breaks the cyclobutane ring joining the two pyrimidines of the dimer. |
| Rhamnogal_lyase | Rhamnogalacturonate lyase ( EC:4.2.2.-) degrades the rhamnogalacturonan I (RG-I) backbone of pectin. This family contains mainly members from plants, but also contains the plant pathogen Erwinia chrysanthemi. | EBP | Emopamil binding protein, encodes a non-glycosylated type I integral membrane protein of endoplasmic reticulum and shows high level expression in epithelial tissues. The EBP protein has emopamil binding domains, including the sterol acceptor site and the catalytic centre, which show Delta8-Delta7 sterol isomerase activity. Human sterol isomerase, a homologue of mouse EBP, is suggested not only to play a role in cholesterol biosynthesis, but also to affect lipoprotein internalisation. |
| Rubrerythrin | This domain has a ferritin-like fold | ETC_C1_NDUFA4 | NADH dehydrogenase (ubiquinone) |

| | | | |
|---|---|---|---|
| rve | Integrase, Retroviral integrase (IN) is an enzyme produced by a retrovirus (such as HIV) that enables its genetic material to be integrated into the DNA of the infected cell. Retroviral INs are not to be confused with phage integrases, such as λ phage integrase (Int) (see site-specific recombination).<br>IN is a key component in the retroviral pre-integration complex (PIC). The complex of integrase bound to cognate viral DNA (vDNA) ends has been referred to as the intasome. | ETF_QO | Electron transfer flavoprotein-ubiquinone oxidoreductase, 4Fe-4S |
| RVT_2 | Reverse transcriptase (RNA-dependent DNA polymerase), is usually indicative of a mobile element such as a retrotransposon or retrovirus. Reverse transcriptases occur in a variety of mobile elements, including retrotransposons, retroviruses, group II introns, bacterial msDNAs, hepadnaviruses, and caulimoviruses. This Pfam entry includes reverse transcriptases not recognised by the PF00078 model. | FAD_binding_7 | Photolyases (EC 4.1.99.3) are DNA repair enzymes that repair damage caused by exposure to ultraviolet light. |
| SASA | Carbohydrate esterase, sialic acid-specific acetylesterase. Sialic acid acetylesterase in autoimmunity | FAD_binding_8 | FAD-binding domain |
| Stealth_CR1 | Stealth protein CR1, Stealth_C1 is the first of several highly conserved regions on stealth proteins in metazoa and bacteria. There are up to four CR regions on all member proteins. CR1 carries a well-conserved IDVVYT sequence-motif. The domain is found in tandem with CR2, CR3 and CR4 on both potential metazoan hosts and pathogenic eubacterial species that are capsular polysaccharide phosphotransferases. The CR domains appear on eukaryotic proteins such as GNPTAB, N-acetylglucosamine-1-phosphotransferase subunits alpha/beta. Horizontal gene-transfer seems to have occurred between host and bacteria of these sequence-regions in order for the bacteria to evade detection by the host innate immune system | Ferric_reduct | Ferric reductase like transmembrane component. This family includes a common region in the transmembrane proteins mammalian cytochrome B-245 heavy chain (gp91-phox), ferric reductase transmembrane component in yeast and respiratory burst oxidase from mouse-ear cress. |
| Stealth_CR2 | Stealth_CR2 is the second of several highly conserved regions on stealth proteins in metazoa and bacteria. There are up to four CR regions on all member proteins. CR2 carries a well-conserved NDD sequence-motif. The domain is found in tandem with CR1, CR3 and | FLILHELTA | protein of unknown function |

| | | | |
|---|---|---|---|
| | CR4 on both potential metazoan hosts and pathogenic eubacterial species that are capsular polysaccharide phosphotransferases. The CR domains appear on eukaryotic proteins such as GNPTAB, N-acetylglucosamine-1-phosphotransferase subunits alpha/beta. Horizontal gene-transfer seems to have occurred between host and bacteria of these sequence-regions in order for the bacteria to evade detection by the host innate immune system | | |
| YoeB_toxin | A toxin-antitoxin system is a set of two or more closely linked genes that together encode both a protein 'poison' and a corresponding 'antidote'. When these systems are contained on plasmids – transferable genetic elements – they ensure that only the daughter cells that inherit the plasmid survive after cell division. If the plasmid is absent in a daughter cell, the unstable antitoxin is degraded and the stable toxic protein kills the new cell; this is known as 'post-segregational killing' (PSK). Toxin-antitoxin systems are widely distributed in prokaryotes, and organisms often have them in multiple copies. | FPN1 | Ferroportin1, that may play a role in iron export from the cell. This family may represent a number of transmembrane regions in Ferroportin1 |
| ZinT | There is strong evidence for involvement of the ZinT domain in zinc homeostasis and management of zinc in the periplasm. It may also facilitate zinc uptake from the environment through interactions with the znuABC zinc transporter. It is regulated by the metalloregulator gene Zur (zinc uptake regulator).<br>The domain was originally discovered in the bacterial stress response to cadmium. Further studies have found that it binds to cadmium, zinc, nickel, and mercury, but not other common metals such as cobalt, copper, iron, and manganese. It may have a secondary function in managing heavy-metal toxicity. | Glyco_hydro_63 | Glycosyl hydrolase family 63 (CAZY GH_63) is a family of eukaryotic enzymes. They catalyse the specific cleavage of the non-reducing terminal glucose residue from Glc(3)Man(9)GlcNAc(2). Mannosyl oligosaccharide glucosidase EC 3.2.1.106 is the first enzyme in the N-linked oligosaccharide processing pathway. |
| | | Glyco_hydro_63N | Glycosyl hydrolase family 63 (CAZY GH_63) is a family of eukaryotic enzymes. They catalyse the specific cleavage of the non-reducing terminal glucose residue from Glc(3)Man(9)GlcNAc(2). Mannosyl oligosaccharide glucosidase EC 3.2.1.106 is the first enzyme in the N-linked oligosaccharide processing pathway |

| | | | |
|---|---|---|---|
| | | Glyco_transf_90 | This family of glycosyl transferases are specifically (mannosyl) glucuronoxylomannan/galactoxylomannan -beta 1,2-xylosyltransferases, EC:2.4.2.-. |
| | | Glyoxal_oxid_N | Glyoxal oxidase N-terminus |
| | | HAGH_C | Hydroxyacylglutathione hydrolase C-terminus. Substrate binding occurs at the interface between this domain and the catalytic domain |
| | | IDO | Indoleamine 2,3-dioxygenase. Indoleamine 2,3-dioxygenase is the first and rate-limiting enzyme of tryptophan catabolism through the kynurenine pathway, thus causing depletion of tryptophan which can cause halted growth of microbes as well as T cells. |
| | | IGR | IGR protein motif |
| | | INSIG | Insulin-induced protein, found in the endoplasmic reticulum and bind the sterol-sensing domain of SREBP cleavage-activating protein (SCAP), preventing it from escorting SREBPs to the Golgi. Their combined action permits feedback regulation of cholesterol synthesis over a wide range of sterol concentrations. |
| | | LIAS_N | LIAS_N is found as the N-terminal domain of the Radical_SAM family in the members that are lipoyl synthase enzymes, particularly the mitochondrial ones in metazoa but also those in bacteria. |
| | | LigB | Catalytic LigB subunit of aromatic ring-opening dioxygenase |
| | | Lipid_DES | Sphingolipid Delta4-desaturase (DES). Sphingolipids are important membrane signalling molecules involved in many different cellular functions in eukaryotes. |
| | | MAM33 | Mitochondrial glycoprotein |
| | | MCD | This family consists of several eukaryotic malonyl-CoA decarboxylase (MLYCD) proteins. Malonyl-CoA, in addition to being an intermediate in the de novo synthesis of fatty acids, is an inhibitor of carnitine palmitoyltransferase I, the enzyme that regulates the transfer of long-chain fatty acyl-CoA into mitochondria, where they are oxidised. |
| | | Mg_trans_NIPA | Magnesium transporter NIPA |

| | | | |
|---|---|---|---|
| | | Mgm101p | Mitochondrial genome maintenance MGM11 |
| | | Mit_KHE1 | Mitochondrial K+-H+ exchange-related |
| | | Mitofilin | Mitochondrial inner membrane protein |
| | | MMADHC | Methylmalonic aciduria and homocystinuria type D protein |
| | | Mo-co_dimer | Mo-co oxidoreductase dimerisation domain. This domain is found in molybdopterin cofactor (Mo-co) oxidoreductases. It is involved in dimer formation, and has an Ig-fold structure |
| | | MOSC_N | MOSC N-terminal beta barrel domain, predicted sulfur-carrier domain |
| | | MRP_L53 | 39S ribosomal protein L53 |
| | | MRP-L20 | Mitochondrial ribosomal protein subunit L2 |
| | | MRP-L28 | Mitochondrial ribosomal protein L28 |
| | | MRP-L46 | 39S mitochondrial ribosomal protein L46 |
| | | MRP-L47 | 39S mitochondrial ribosomal protein L47 |
| | | MRP-S25 | Mitochondrial ribosomal protein S25 |
| | | MTP18 | Mitochondrial 18 KDa protein |
| | | MWFE | NADH dehydrogenase (ubiquinone), is an enzyme of the respiratory chains of myriad organisms from bacteria to humans that falls under the H+ or Na+-translocating NADH Dehydrogenase (NDH) Family (TC# 3.D.1), a member of the Na+ transporting Mrp superfamily. |
| | | NAD_binding_6 | Ferric reductase NAD binding domain Provide feedback |
| | | NADH-u_ox-rdase | NADH-ubiquinone oxidoreductase complex I, 21 kDa subunit |
| | | NDUF_B7 | NADH dehydrogenase (ubiquinone) |
| | | NTPase_I-T | protein of unknown function |
| | | Ofd1_CTDD | Oxoglutarate and iron-dependent oxygenase degradation C-term |
| | | OHCU_decarbox | In molecular biology 2-oxo-4-hydroxy-4-carboxy-5-ureidoimidazoline decarboxylase (OHCU decarboxylase) EC 4.1.1.n1 is an enzyme involved in purine catabolism |
| | | OPA3 | Optic atrophy 3 protein, deficiency of which causes type III 3-methylglutaconic aciduria (MGA) in humans. This disease manifests with early bilateral optic atrophy, spasticity, extrapyramidal |

| | | | |
|---|---|---|---|
| | | | dysfunction, ataxia, and cognitive deficits, but normal longevity |
| | | Pam17 | Mitochondrial import protein Pam17 |
| | | PDZ_1 | These domains play a key role in the formation and function of signal transduction complexes. |
| | | Peptidase_M76 | This is a family of metalloproteases. Proteins in this family are also annotated as Ku70-binding proteins. |
| | | PET117 | PET assembly of cytochrome c oxidase |
| | | Pet127 | Mitochondrial protein Pet127 |
| | | Pet191_N | Cytochrome c oxidase |
| | | Pirin_C | Pirin C-terminal cupin domain |
| | | QCR10 | Ubiquinol-cytochrome-c reductase complex subunit |
| | | Rad52_Rad22 | The DNA single-strand annealing proteins (SSAPs), such as RecT, Red-beta, ERF and Rad52, function in RecA-dependent and RecA-independent DNA recombination pathways |
| | | RdRP | RNA dependent RNA polymerase, This family of proteins are eukaryotic RNA dependent RNA polymerases. These proteins are involved in post transcriptional gene silencing where they are thought to amplify dsRNA templates. |
| | | Rib_5-P_isom_A | Ribose 5-phosphate isomerase A (phosphoriboisomerase A) |
| | | Ribosomal_L33 | unknown |
| | | Ribosomal_L34 | unknown |
| | | Ribosomal_L36 | unknown |
| | | RPOL_N | DNA-directed RNA polymerase N-terminal. This domain has a role in interaction with regions of upstream promoter DNA and the nascent RNA chain, leading to the processivity of the enzyme. This domain undergoes a structural change in the transition from initiation to elongation phase. The structural change results in abolition of the promoter binding site, creation of a channel accommodating the heteroduplex in the active site and formation of an exit tunnel which the RNA transcript passes through after peeling off the heteroduplex. |
| | | S1-P1_nuclease | S1 P1 nuclease protein domain, which cleave RNA and single stranded DNA with no sequence specificity. They are |

| | | | |
|---|---|---|---|
| | | | found in both prokaryotes and eukaryotes and are thought to be associated in programmed cell death and also in tissue differentiation. Furthermore, they are secreted extracellular, that is, outside of the cell. Their function and distinguishing features mean they have potential in being exploited in the field of biotechnology. |
| | | SCO1-SenC | This family is involved in biogenesis of respiratory and photosynthetic systems. |
| | | SE | Squalene epoxidase. This domain is found in squalene epoxidase (SE) and related proteins which are found in taxonomically diverse groups of eukaryotes and also in bacteria. SE was first cloned from Saccharomyces cerevisiae where it was named ERG1. It contains a putative FAD binding site and is a key enzyme in the sterol biosynthetic pathway [1]. Putative transmembrane regions are found to the protein's C-terminus. |
| | | Sgf11 | The Sgf11 family is a SAGA complex subunit in Saccharomyces cerevisiae. The SAGA complex is a multisubunit protein complex involved in transcriptional regulation. SAGA combines proteins involved in interactions with DNA-bound activators and TATA-binding protein (TBP), as well as enzymes for histone acetylation and deubiquitylation |
| | | She9_MDM33 | mitochondrial inner membrane proteins with a role in inner mitochondrial membrane organisation and biogenesis |
| | | Solute_trans_a | Organic solute transporter Ostalpha. This family is a transmembrane organic solute transport protein. In vertebrates these proteins form a complex with Ostbeta, and function as bile transporters [1]. In plants they may transport brassinosteroid-like compounds and act as regulators of cell death |
| | | Spo12 | The Spo12 protein plays a regulatory role in two of the most fundamental processes of biology, mitosis and meiosis, and yet its biochemical function remains elusive. Spo12 is a nuclear protein. Spo12 is a component of the FEAR (Cdc fourteen early anaphase release) regulatory network, that promotes Cdc14 release from the nucleolus during early anaphase. The FEAR network is comprised of the polo kinase Cdc5, the |

| | | | |
|---|---|---|---|
| | | | separase Esp1, the kinetochore-associated protein Slk19, and Spo12. |
| | | SPO22 | SPO22/ZIP4 in yeast is a meiosis specific protein involved in sporulation. It has been shown to regulate crossover distribution by promoting synaptonemal complex formation |
| | | Suc_Fer-like | Sucrase/ferredoxin-like. This family contains a number of bacterial and eukaryotic proteins approximately 400 residues long that resemble ferredoxin and appear to have sucrolytic activity |
| | | TIM21 | TIM21 interacts with the outer mitochondrial TOM complex and promotes the insertion of proteins into the inner mitochondrial membrane |
| | | TMEM223 | Transmembrane protein 223 |
| | | UcrQ | Coenzyme Q – cytochrome c reductase |
| | | Ureidogly_lyase | Ureidoglycolate lyase, one of the enzymes that acts upon ureidoglycolate, an intermediate of purine catabolism, releasing urea. |
| | | VRR_NUC | unknown |
| | | YCII | YCII-related domain. This domain is suggested to play a role in transcription initiation (Bateman A per. obs.). This domain is named after the most conserved motif in the alignment. |
| | | zf-4CXXC_R1 | Zinc-finger domain of monoamine-oxidase A repressor R1 |
| | | zf-CHCC | NADH dehydrogenase (ubiquinone) |

**Table S2.** Genome information of the animal hosts and diet plants used in the study to infer the genetic elements in Neocallimastigomycota that have a foreign origin

| Name | Taxon | Version | Source |
|---|---|---|---|
| Elephant | *Loxodonta africana* (African savanna elephant) | Loxafr3.0 | Broad Institute |
| Horse | *Equus caballus* | EquCab2.0 | Wade et al. 2009 |
| Sheep | *Ovis aries* | Oar_v4.0 | International Sheep Genomics Consortium 2010 |
| Yak | *Bos mutus* (wild yak) | BosGru_v2.0 | Qiu et al. 2012 |
| Banana | *Musa acuminata* | DH-Pahang v2 | Martin et al. 2016 |
| Palm | *Elaeis guineensis* (African oil palm) | EG5 | Singh et al. 2013 |
| Bamboo | *Phyllostachys heterocycla* var. pubescens | v1.0 | Peng et al. 2013 |
| GoatGrass | *Aegilops tauschii* subsp. Tauschii | Aet_MR_1.0 | Zimin et al. 2017 |
| Maize | *Zea mays* | B73 RefGen_v4 | Jiao Y, et al. 2017 |
| Rice | *Oryza sativa* Japonica Group | Build 4.0 | Rice Annotation Project 2007 |
| Brome | *Brachypodium distachyon* | v2.0 | International Brachypodium Initiative 2010 |
| Sorghum | *Sorghum bicolor* | v3 | Paterson et al. 2009 |
| Arabidopsis | *Arabidopsis thaliana* | TAIR10 | Swarbreck et al. 2008 |
| Moss | *Physcomitrella patens* | V1.1 | Rensing et al. 2008 |
