## Supplementary results and associated figures for "Molecular dating of the emergence of anaerobic rumen fungi and the impact of laterally acquired genes"

**Supplementary results for putative intron gain in AGF "Cthe_2159" genes**

Approximately 30% of the "Cthe_2159" genes across all publicly available AGF genomes possess 1-2 introns according to the genome annotation (Grigoriev et al., 2014; Haitjema et al., 2017; Youssef et al., 2013). Phylogenetic analysis demonstrated that "Cthe_2159" genes locating on the same scaffold tend to cluster with each other with greater than 99/100 bootstrap supports, regardless of the intron numbers (0-2) they possess (Supplementary Figure A-a). This suggests that the pattern of intron presence in AGF "Cthe_2159" is random without a major constraint evolutionary event above genus level. Some introns, especially in *Piromyces* sp. E2 are adjacent to duplicated fragments of coding sequences within the conserved "Cthe_2159" domain (e.g., PirE2_1_10137 in Supplementary Figure A-b and PirE2_1_5444 in Supplementary Figure A-c). The duplication is apparent by comparing alignments of the nucleotide sequences (Supplementary Figure A-b and A-c). We hypothesize these introns represent a gain as a result of a local duplication event according to the proposed "Tandem Genomic Duplication" model of intron gains (Yenerall and Zhou, 2012). Unfortunately, *Piromyces* sp. E2 strain lacks of transcriptome data, which could be used to confirm the reality of reported introns. Although Expressed Sequence Tag (EST) reads are available at JGI website, they are not sufficient to recover the interested regions (PirE2_1_10137 and PirE2_1_5444). In addition, the *Piromyces* sp. E2 genome is a pioneer assembly across the entire lineage (assembled in March 2011) using Sanger gDNA and shredded Velvet contigs (Grigoriev et al., 2014; Haitjema et al., 2017). We cannot exclude the possibility that the annotated introns and flanked duplications were due to assembly artifact, which pends future investigation using better quality genome assemblies if the strain is still alive and accessible.

The genome of *Piromyces finnis* represents the best quality of all AGF genomes with multiple transcriptomic characterization studies (Haitjema et al., 2017; Solomon et al., 2016). To seek evidence to prove the reality of annotated introns in "Cthe_2159" genes, we used splicing-site sensitive tool (Hisat2) to map the transcriptome reads to the *Piromyces finnis* genome assembly. At the interested regions where introns were reported in *Piromyces finnis*, we did not find supportive splicing evidence by examining the paired reads using Integrative Genomics Viewer (IGV) manually (Supplementary Figures B and C) (Robinson et al., 2011). Although the negative evidence found in *Piromyces finnis* cannot be used to explain a crossed strain (*Piromyces* sp. E2) directly, especially considering that the reported introns in *Piromyces* sp. E2 are flanked by duplicated coding sequences, which is usually a strong sign for the intron insertion (Yenerall and Zhou, 2012), the inconsistency between gene annotation and transcriptome evidence in *Piromyces finnis* alert us the intron presence in other AGF genomes may due to similar artifacts.

**Supplementary Methods**

Introns in AGF "Cthe_2159" genes were identified by examining the genome annotation files (Grigoriev et al., 2014; Haitjema et al., 2017; Youssef et al., 2013). The "Cthe_2159" domain containing genes in the 5 AGF genomes were aligned using MUSCLE v3.8.31 in Mesquite software (Edgar, 2004; Maddison and Maddison, 2007). Dot plots to visualize the duplicated coding sequences within the conserved "Cthe_2159" domains were produced using the Dottup tool in EMBOSS v6.5.7 (Rice et al., 2000). Transcriptome reads of *Piromyces finnis* (SRR5487626) were retrieved using the SRA toolkit (Leinonen et al., 2011) and mapped to the genome assembly using Hisat2 (Kim et al., 2015). Regions of interest containing predicted introns were examined manually with IGV (Robinson et al., 2011).

**Supplementary Figure A.** Intron insertion events identified in the “Cthe_2159” genes in the five AGF genomes. (a) Phylogenetic tree of the 83 AGF “Cthe_2159” domains based on protein sequences (rooted with the closest related bacterial homolog). Both intron numbers and domain maps are shown and associated with the tree tips. Highlighted clades are provided with detailed comparative analyses for intron insertions (yellow clade is shown in the part b; green clades are unfolded in the part c). (b) Three highly identical “Cthe_2159” domains (PirE2_1_10132, PirE2_1_10136, and PirE2_1_10137) with various intron numbers (0-2) located on the same scaffold (scaffold_146) of the *Piromyces* sp. E2 genome. Inserted introns can be either within or outside of the “Cthe_2159” domains. Pieces of duplicated regions are found in CDS2 and adjacently connected with the intron, thus likely associated with the intron insertion event in the PirE2_1_10137 gene. This is confirmed by multiple sequence alignments (with both protein and nucleotide level) and self dot-plot analyses. (c) Three pairs of “Cthe_2159” domains are found identical and flanked on a pair of homologous scaffolds in *Piromyces finnis* (scaffold_6) and *Piromyces* sp. E2 (scaffold_46) genomes separately. Multiple intron gain and loss events can be identified by comparing each of the three pairs. A recent gene duplication event (PirE2_1_5444 is a recent duplicate of the PirE2_1_5440 according to the phylogenetic tree in part a) followed by intron insertion (in PirE2_1_5444) is also found with redundant coding sequences within the “Cthe_2159” domain. Similar to the part b, a regional duplication associated with intron insertion is confirmed by both multiple sequence alignment and self dot-plot analyses.

**Supplementary Figure B.** Transcriptome mapping result of the *Piromyces finnis* showing the region of Pirfi3_345772 (both annotated introns were not supported by the transcriptome data).

**Supplementary Figure C.** Transcriptome mapping result of the *Piromyces finnis* showing the region of Pirfi3_414173 (both annotated introns were not supported by the transcriptome data).

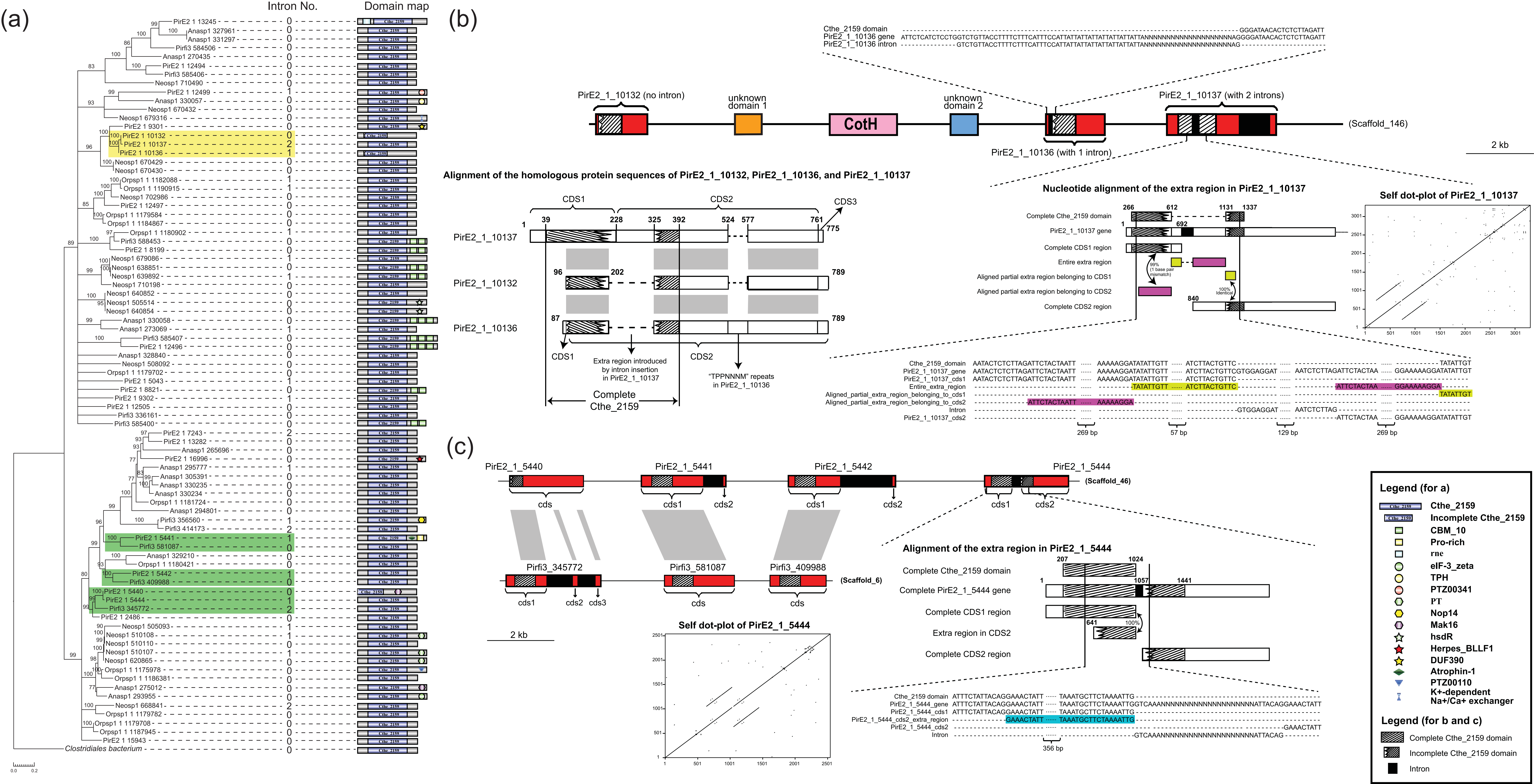

(a)
Intron No.
Domain map
PirE2 1 13245
Anasp1 327961
Anasp1 331297
Pirfi3 584506
Anasp1 270435
PirE2 1 12494
Pirfi3 585406
Neosp1 710490
PirE2 1 12499
Anasp1 330057
Neosp1 670432
Neosp1 679316
PirE2 1 9301
PirE2 1 10132
PirE2 1 10137
PirE2 1 10136
Neosp1 670429
Neosp1 670430
Orpsp1 1 1182088
Orpsp1 1 1190915
Neosp1 702986
PirE2 1 12497
Orpsp1 1 1179584
Orpsp1 1 1184867
Orpsp1 1 1180902
Pirfi3 588453
PirE2 1 8199
Neosp1 679086
Neosp1 638851
Neosp1 639892
Neosp1 710198
Neosp1 640852
Neosp1 505514
Neosp1 640854
Anasp1 330058
Anasp1 273069
Pirfi3 585407
PirE2 1 12496
Anasp1 328840
Neosp1 508092
Orpsp1 1 1179702
PirE2 1 5043
PirE2 1 8821
PirE2 1 9302
PirE2 1 12505
Pirfi3 336161
Pirfi3 585400
PirE2 1 7243
PirE2 1 13282
Anasp1 265696
PirE2 1 16996
Anasp1 295777
Anasp1 305391
Anasp1 330235
Anasp1 330234
Orpsp1 1 1181724
Anasp1 294801
Pirfi3 356560
Pirfi3 414173
PirE2 1 5441
Pirfi3 581087
Anasp1 329210
Orpsp1 1 1180421
PirE2 1 5442
Pirfi3 409988
PirE2 1 5440
PirE2 1 5444
Pirfi3 345772
PirE2 1 2486
Neosp1 505093
Neosp1 510108
Neosp1 510110
Neosp1 510107
Neosp1 620865
Orpsp1 1 1175978
Orpsp1 1 1186381
Anasp1 275012
Anasp1 293955
Neosp1 668841
Orpsp1 1 1179782
Orpsp1 1 1179708
Orpsp1 1 1187945
PirE2 1 15943
Clostridiales bacterium
0.0 0.2
(b)
Cthe_2159 domain
PirE2_1_10136 gene
PirE2_1_10136 intron
PirE2_1_10132 (no intron)
unknown domain 1
CotH
unknown domain 2
PirE2_1_10137 (with 2 introns)
(Scaffold_146)
PirE2_1_10136 (with 1 intron)
2 kb
Alignment of the homologous protein sequences of PirE2_1_10132, PirE2_1_10136, and PirE2_1_10137
CDS1
CDS2
CDS3
PirE2_1_10137
PirE2_1_10132
PirE2_1_10136
Extra region introduced by intron insertion in PirE2_1_10137
"TPPNNNM" repeats in PirE2_1_10136
Complete Cthe_2159
Nucleotide alignment of the extra region in PirE2_1_10137
Complete Cthe_2159 domain
PirE2_1_10137 gene
Complete CDS1 region
Entire extra region
Aligned partial extra region belonging to CDS1
Aligned partial extra region belonging to CDS2
Complete CDS2 region
99% (1 base pair mismatch)
100% Identical
Self dot-plot of PirE2_1_10137
Cthe_2159_domain
PirE2_1_10137_gene
PirE2_1_10137_cds1
Entire_extra_region
Aligned_partial_extra_region_belonging_to_cds1
Aligned_partial_extra_region_belonging_to_cds2
Intron
PirE2_1_10137_cds2
269 bp
57 bp
129 bp
269 bp
(c)
PirE2_1_5440
PirE2_1_5441
PirE2_1_5442
PirE2_1_5444
(Scaffold_46)
cds
cds1
cds2
Pirfi3_345772
Pirfi3_581087
Pirfi3_409988
(Scaffold_6)
cds3
2 kb
Self dot-plot of PirE2_1_5444
Alignment of the extra region in PirE2_1_5444
Complete Cthe_2159 domain
Complete PirE2_1_5444 gene
Complete CDS1 region
Extra region in CDS2
Complete CDS2 region
Cthe_2159 domain
PirE2_1_5444_gene
PirE2_1_5444_cds1
PirE2_1_5444_cds2_extra_region
PirE2_1_5444_cds2
Intron
356 bp
Legend (for a)
Cthe_2159
Incomplete Cthe_2159
CBM_10
Pro-rich
rne
eIF-3_zeta
TPH
PTZ00341
PT
Nop14
Mak16
hsdR
Herpes_BLLF1
DUF390
Atrophin-1
PTZ00110
K+-dependent Na+/Ca+ exchanger
Legend (for b and c)
Complete Cthe_2159 domain
Incomplete Cthe_2159 domain
Intron

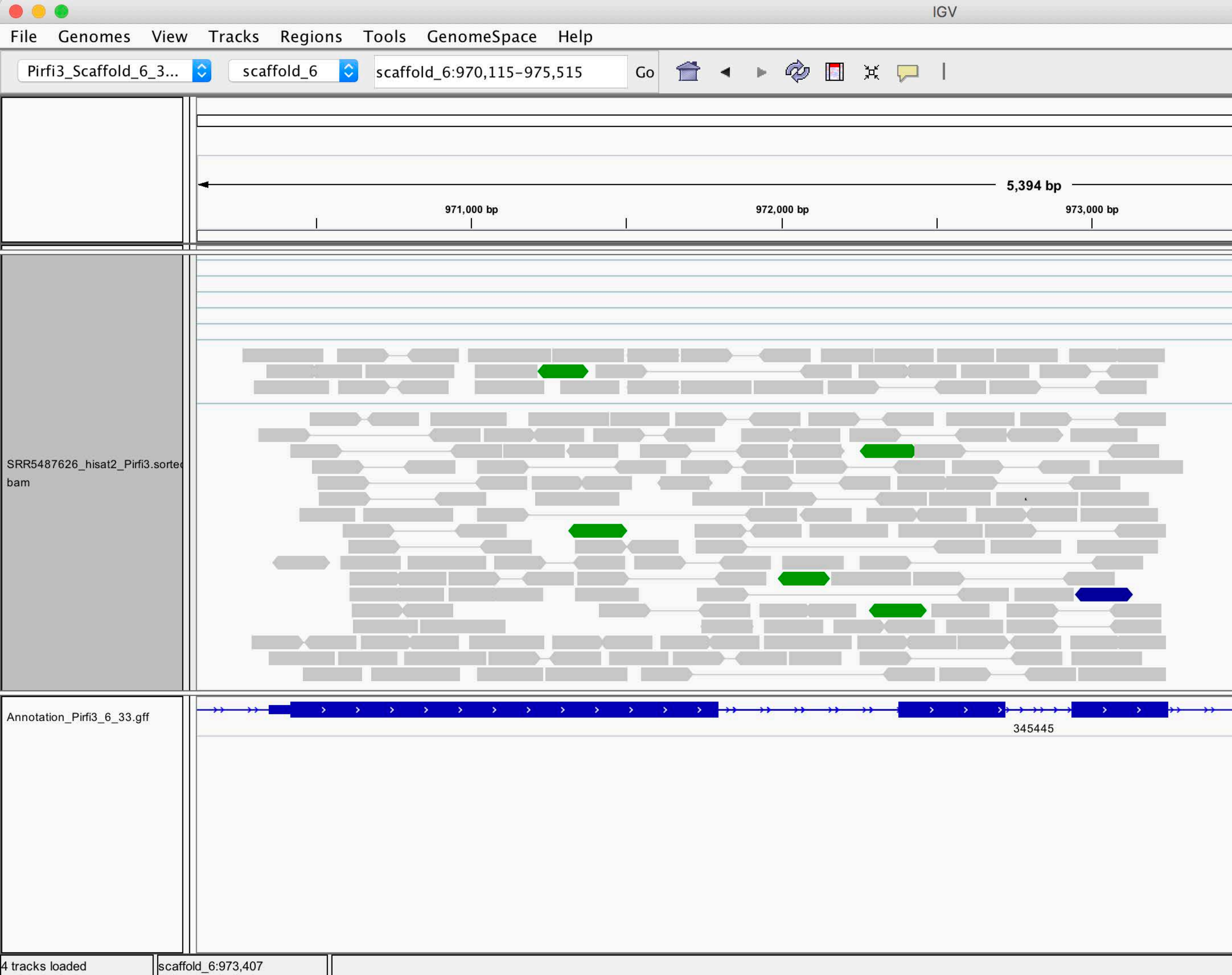
IGV
File
Genomes
View
Tracks
Regions
Tools
GenomeSpace
Help
Pirfi3_Scaffold_6_3...
scaffold_6
scaffold_6:970,115-975,515
Go
5,394 bp
971,000 bp
972,000 bp
973,000 bp
SRR5487626_hisat2_Pirfi3.sorted
bam
Annotation_Pirfi3_6_33.gff
345445
4 tracks loaded
scaffold_6:973,407

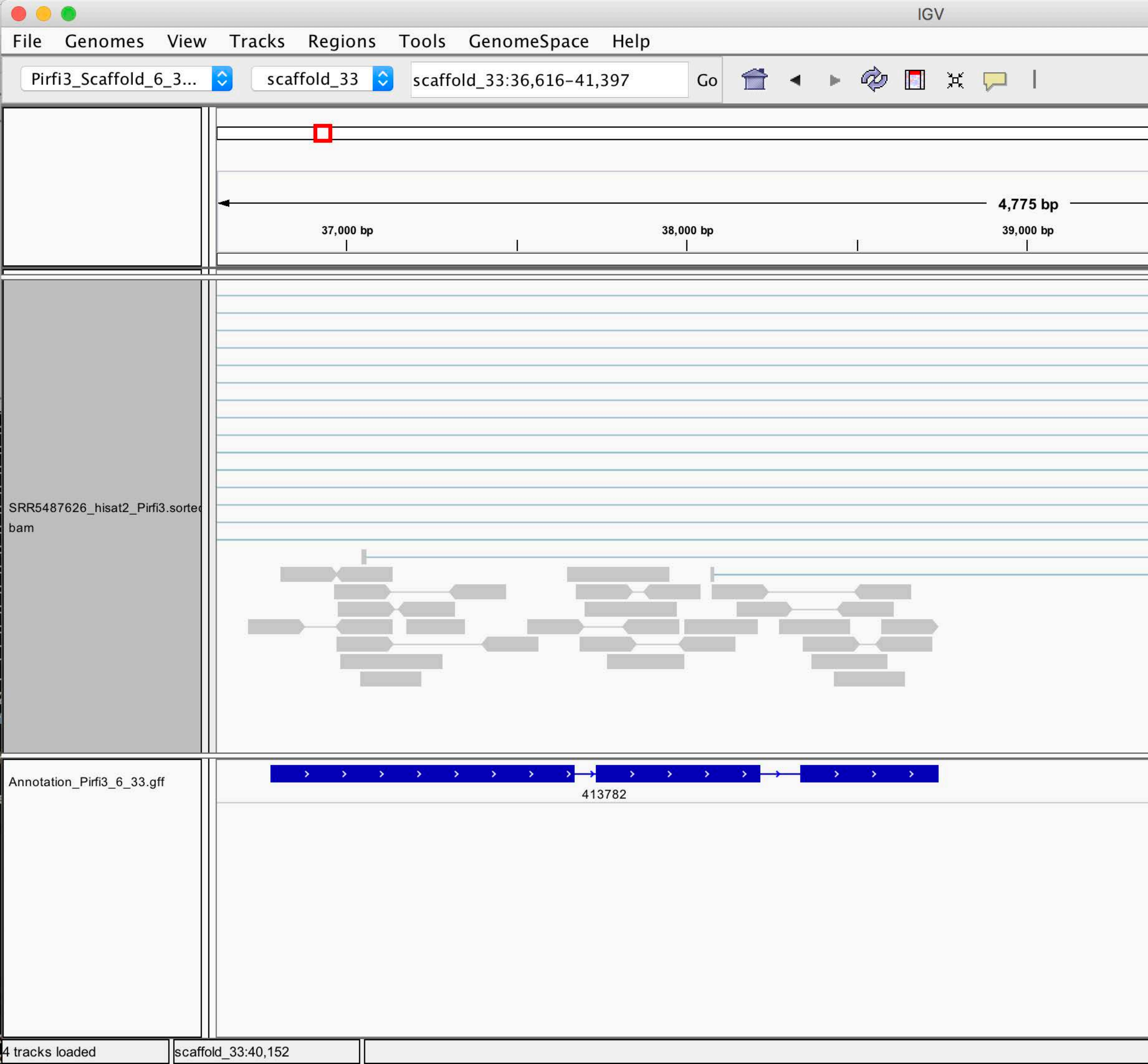
IGV
File Genomes View Tracks Regions Tools GenomeSpace Help
Pirfi3_Scaffold_6_3...
scaffold_33
scaffold_33:36,616-41,397
Go
4,775 bp
37,000 bp
38,000 bp
39,000 bp
SRR5487626_hisat2_Pirfi3.sorted.bam
Annotation_Pirfi3_6_33.gff
413782
4 tracks loaded
scaffold_33:40,152
